# Reactivation of a TAL1 progenitor cell enhancer region by non-coding somatic variants in T-lineage acute lymphoblastic leukemia

**DOI:** 10.64898/2026.05.03.722504

**Authors:** Nadezhda V. Terekhanova, Xiaolong Chen, Kin-Hoe Chow, Yu Liu, Ying Shao, Li Dong, Bensheng Ju, Vinesh Vinayachandran, Haseeb Zubair, Kohei Hagiwara, Wentao Yang, Xiaotu Ma, Sivaraman Natarajan, John Easton, David T. Teachey, A. Thomas Look, Jinghui Zhang

**Affiliations:** Department of Computational Biology, St. Jude Children’s Research Hospital, Memphis, TN, 38105, USA; Pediatric Translational Medicine Institute, Shanghai Children’s Medical Center, School of Medicine, Shanghai Jiao Tong University, Shanghai, China; Department of Pediatric Oncology, Dana-Farber Cancer Institute, Harvard Medical School, Boston, MA, 02215, USA; Perelman School of Medicine, University of Pennsylvania, Philadelphia, PA, 19104, USA; Division of Oncology and Center for Childhood Cancer Research, Children’s Hospital of Philadelphia, Philadelphia, PA, 19104, USA; Department of Pediatrics and the Center for Childhood Cancer Research, Children’s Hospital of Philadelphia, Philadelphia, PA, 19104, USA; Division of Pediatric Hematology and Oncology, Boston Children’s Hospital, Boston, MA, 02115, USA

## Abstract

Aberrant activation of *TAL1*, a key oncogenic driver, defines a major subgroup comprising ∼30% of childhood T-lineage acute lymphoblastic leukemias (T-ALLs). We and others have shown that somatic non-coding mutations within upstream and intronic *cis*-regulatory regions of *TAL1* contribute to transformation by creating binding sites for MYB and other transcription factors. Here we investigated *cis*-regulatory mechanisms mediated by somatic mutations occurring in an intergenic region located 29 kilobase pairs downstream of the canonical *TAL1* transcription initiation site, implicated in 6% of *TAL1*-expressing T-ALLs. These somatic variants include i) complex indels resulting in *de novo* MYB transcription factor binding sites (TFBSs) and ii) internal tandem duplications (ITDs) encompassing canonical MYB TFBSs. Chromatin immunoprecipitation sequencing (ChIP-seq) revealed binding of the TAL1 core regulatory circuit (CRC) transcription factors MYB, GATA3, and RUNX1, resulting in enhancer activity mediated by sequences with the mutant allele. Strikingly, ChIP-seq peaks for the repressive H3K27me3 mark and the active H3K27ac mark co-existed across *TAL1* regulatory sequences but enriched for different haplotypes. *TAL1* transcription from the mutant haplotype initiated from a promoter located within exon 4 of the canonical *TAL1* transcript, resulting in a short isoform normally expressed by hematopoietic stem cells (HSC). Interestingly, neither the isoform expression nor the enhancer activity could be predicted by the sequence-to-function deep learning artificial intelligence (AI) model AlphaGenome, emphasizing the importance of experimental validation. Our findings indicate that selection for *cis*-regulatory, non-coding variants leads to reactivation of enhancers normally active in HSC but silenced in differentiated lineages during normal hematopoietic cell development.

## Introduction

*TAL1*, also known as *SCL*, is an oncogene encoding a basic helix-loop-helix (bHLH) transcription factor, which functions as a key driver in T-lineage acute lymphoblastic leukemia (T-ALL). Also known as a critical regulator for normal hematopoietic cell development, *TAL1* is essential for primitive hematopoiesis during embryogenesis^1,2^, as well as the formation of hematopoietic stem cells (HSCs) and lineage commitment during erythroid and megakaryocytic cell development throughout life^3,4^. *TAL1* is expressed by HSCs, multipotent progenitors (MPPs), and erythroid and megakaryocytic lineages, but silenced during B- and T-cell development^5,6^. Lineage-specific *TAL1* expression is controlled by different enhancers, as identified by epigenetic profiling as well as by transgenic reporter and rescue experiments. There are three well-studied enhancers, which include two downstream regions named according to their distance in kilobase pairs (kbp) from the *TAL1* promoter, i) the +51 erythroid enhancer^7^, ii) the +19/20/21 hematopoietic progenitor cell enhancer^8^ (also known as the stem cell enhancer^9^), and iii) 5’ enhancer located 4 kbp upstream of the promoter, which also regulates the expression in normal progenitor cells^10^.

In approximately 30-40% of the T-ALLs, aberrant activation of *TAL1* defines a distinct subtype *TAL1* T-ALL^11,12^. This is achieved by somatically acquired genomic re-arrangements, or mutations such as single nucleotide variations (SNVs) or small insertion/deletions (indels) in the non-coding regions, which mediate *cis*-regulation of *TAL1* transcription. The chromosomal re-arrangements include translocations resulting in juxtaposition of T-cell receptor (TCR) *cis*-regulatory elements adjacent to the *TAL1* gene^13^; and an intergenic deletion between the *STIL* and *TAL1* genes on the short arm of human chromosome 1, which places the *TAL1* coding sequences under the regulatory control of the adjacent *STIL* gene regulatory sequences^14^. In addition, *cis*-regulatory mutations affect multiple non-coding regions of the *TAL1* locus as shown in **Fig. 1a**. The first to be described is a recurrent insertion at the hotspot located 7 kbp upstream of *TAL1* coding sequences, leading to the creation of *de novo* MYB transcription factor binding site (TFBS), which allows MYB binding and opens adjacent chromatin, facilitating transcription factors of the TAL1 Core Regulatory Circuit (CRC), such as

**Fig. 1:**
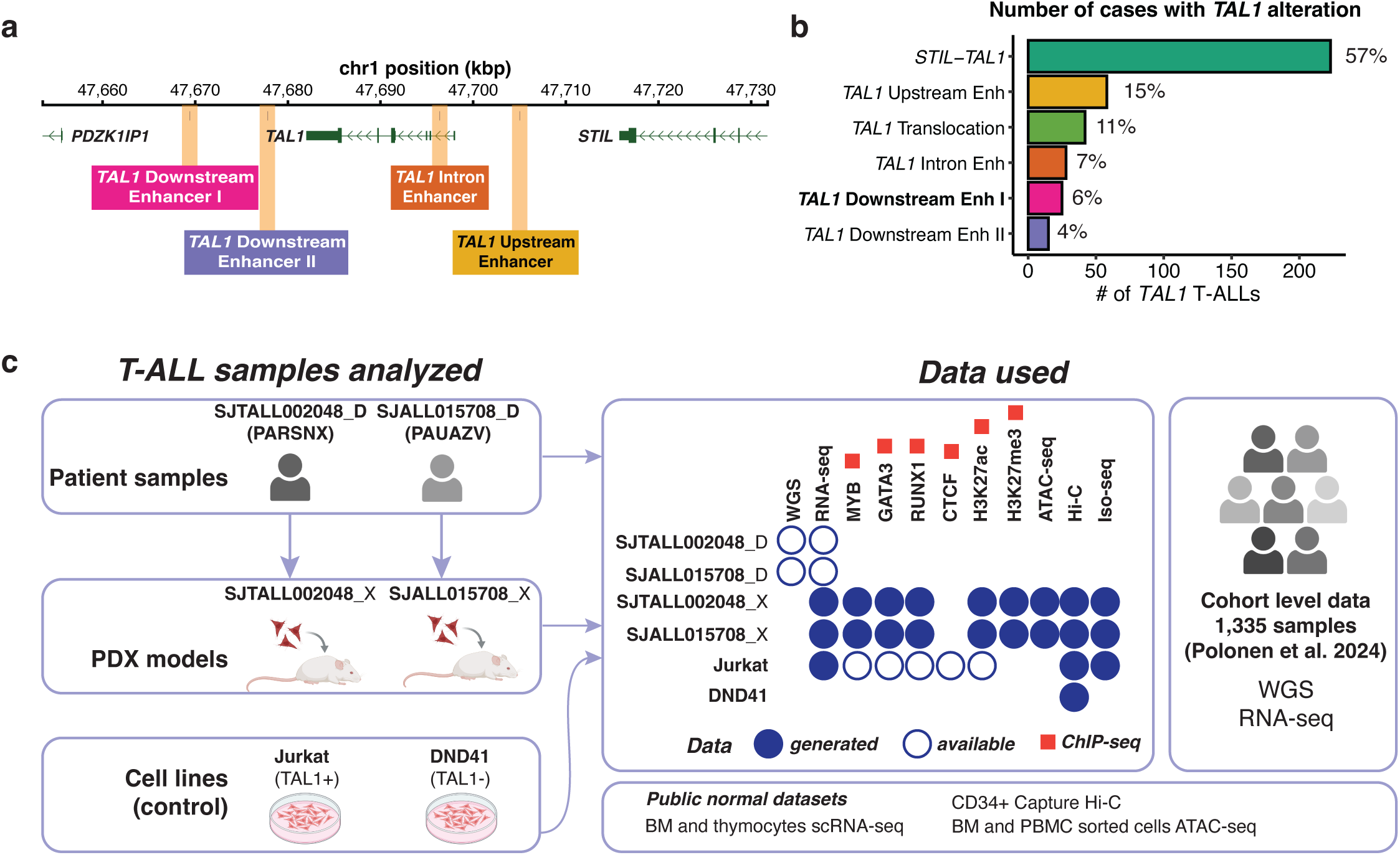
**Overview of *TAL1* alterations in T-ALL and study design for somatic variants at downstream enhancer I region**. **a**, Genomic view showing the locations of four *TAL1* enhancers caused by non-coding variants. **b**, Barplot showing the distribution of T-ALL samples across six groups of somatic alterations leading to *TAL1* activation. At the right side of each bar, the percentage relative to the total number of samples with *TAL1* alterations is shown. The “TAL1 Downstream Enh I” group shown in bold is the focus of the current study. **c**, Schematic showing the two index samples from “TAL1 Downstream Enh I” group from which the PDX models were developed for multi-omics profiling. Additionally public data sets generated from patient T-ALL samples and normal hematopoietic development were also used. BM corresponds to bone marrow and PBMC corresponds to peripheral blood mononuclear cells. Created with BioRender.com.

GATA3 and RUNX1, to bind their consensus sequences, which together mediate the formation of an active super enhancer^15^. A second group of *TAL1 cis*-regulatory mutations in T-ALL occurs in the form of a recurrent SNV in *TAL1* intron 1, creating a *de novo* YY1 TFBS associated with YY1 binding and elevated enhancer activity^16^. In a third group of *TAL1* T-ALLs, recurrent insertions are found 20 kbp downstream of the *TAL1* gene, leading to the creation of MYB TFBSs and the formation of a super-enhancer^17^. Finally, in a fourth group of *TAL1* T-ALLs, a series of somatic indels, inversions and internal tandem duplications (ITDs) occur, located approximately 29 kbp downstream of *TAL1*^12^. In a recent analysis of 1,309 T-ALL whole genome sequencing (WGS) samples by Polonen et al.^12^, the somatic variants in these four non-coding regions account for a significant proportion of *TAL1* T-ALLs (**Fig. 1b**) and are denoted as i) *TAL1* upstream enhancer, ii) *TAL1* intron enhancer, iii) *TAL1* downstream enhancer II and iv) *TAL1* downstream enhancer I.

The MYB transcription factor (TF) has been shown to be required during embryogenesis for the normal development of both T and B lymphocytes^18^. Within the T-cell lineage, MYB is expressed at high levels by both double-negative and CD4+/CD8+ double-positive thymocytes, and then MYB is down-regulated as the thymocytes undergo TCR alpha chain rearrangement, which is followed by positive and negative selection and commitment to become either CD4 or CD8 single-positive thymocytes. Selection for *de novo* mutations that form MYB binding sites during T-ALL transformation imply that MYB is expressed by the thymocyte progenitors that are the cells of origin for these leukemias, suggesting that they arise from CD4/CD8 double-negative or double-positive thymocytes^19^. Mutations in the upstream 5’ enhancer^15^ and the majority of mutations in 3’ downstream enhancer II region of *TAL1* occur as small indels^17^ resulting in *de novo* MYB TFBSs, and enhancer activation upon MYB binding at these sites has been validated by chromatin immunoprecipitation sequencing (ChIP-seq) analysis. MYB is also known to be a pioneer factor that can bind even condensed, “closed” chromatin and initiate its opening and then recruit other core TFs that together form an active super-enhancer driving high levels of *TAL1* expression^20,21^. This is consistent with the observation in human T-ALLs that after MYB binds to its new mutant binding site, the other transcription factors of the TAL1 CRC, such as GATA3 and RUNX1, are able to bind to their recognition sequences adjacent to MYB. Furthermore, this observation emphasizes that the *de novo* somatic MYB TFBS location is constrained by requiring the nearby presence of TFBSs of members of TAL1 CRC, i.e. RUNX1, GATA3, ETS and E-box proteins^15,22^.

While recurrent somatic variants mediating the super enhancers that form in upstream enhancer, intron 1 and downstream enhancer II were shown in prior studies to have enhancer activity leading to mono-allelic expression of *TAL1*, there is limited knowledge about the regulatory mechanisms mediated by diverse variants in the non-coding region that form downstream enhancer I. It is known that these variants lead to open chromatin, measured by assay for transposase-accessible chromatin using sequencing (ATAC-seq), and also to the formation of a large super enhancer with extensive H3K27ac chromatin modifications in at least one sample^12^. Although all non-coding variants upstream or in intron 1 have been considered to generate neo-enhancers that are not found in normal cells, this distinction from normal *TAL1* enhancer regions remains uncertain for the less well-studied *TAL1* downstream enhancers, because of their physical proximity to the +19/20/21 stem cell enhancer active in normal hematopoietic progenitor cells.

Previously, when investigating *TAL1* T-ALLs that lacked canonical re-arrangements but exhibited elevated and mono-allelic *TAL1* expression, we detected a complex somatic indel and an ITD in two patient T-ALLs, SJTALL002048_D and SJALL015708_D, respectively, within downstream enhancer I region. Patient-derived xenograft (PDX) mouse models were developed from these two T-ALLs, which immortalized these cells and allowed us to investigate *cis*-regulatory mechanisms by performing epigenetic profiling, 3D genome mapping, and full-length transcriptome sequencing (**Fig. 1c**). As controls for this investigation, we also used multi-omic data from two T-ALL cell lines Jurkat (a *TAL1* positive cell line with an upstream enhancer indel^15^) and DND41 (a *TAL1* negative cell line), which were either studied experimentally or obtained from public repositories. Our investigation involved integration of T-ALL epigenetic data with multiple public datasets that had been generated from different cell types during normal hematopoietic cell development (**Fig. 1c**). These analyses led to the new insights into *TAL1* enhancer architecture in T-ALL, regulation of *TAL1* long-vs-short isoform expression, and associations with T-cell developmental stages at cancer initiation, which involves the reactivation of an enhancer which is normally silenced during thymocyte differentiation into single-positive mature T cells.

In this study, we also performed a comparison of our findings derived from PDX models with *in silico* prediction by the recently published AlphaGenome^23^, a deep-learning artificial intelligence (AI) model for sequence-to-function analysis. In the AlphaGenome study, *TAL1* non-coding variants reported in the prior published enhancer regions (i.e. upstream enhancer^15^, intron enhancer^16^, and downstream enhancer II^17^) were tested to highlight AlphaGenome’s accuracy on regulatory variant effect prediction. By contrast, variants in *TAL1* downstream enhancer I studied here were not analyzed, which allowed us to perform an unbiased evaluation on the performance of AlphaGenome. Our results demonstrate the importance of generating experimental data to gain insight into novel regulatory mechanisms.

## Results

### Overlap of *TAL1* somatic non-coding variants in T-ALL with enhancers in hematopoietic progenitor cells

Aberrant activation of *TAL1* in T-ALL can occur through mutational alterations within four separate non-coding regions (**Fig. 1a**), by *STIL-TAL1* deletion or *TAL1* translocations. To ensure an accurate assessment on the prevalence of these events and precise mapping of their genomic locations, we first curated the 404 T-ALLs denoted to have somatic alterations associated with *TAL1* activation by Polonen et al.^12^, which also included our two index cases (see Methods and **Supplementary Table 1**). Four samples were re-assigned as their variant locations matched a different *TAL1* non-coding enhancer region and 13 were excluded due to: i) presence of variants in multiple regions (n=5); ii) mismatch of variant locations with all denoted enhancer regions (n=5); and iii) SVs corresponding to *TAL1* inversions involving *TAL1* exons and introns instead of the canonical *TAL1* translocation events (n=3). Using the remaining 391 curated T-ALLs, we determined the prevalence of these six types of *TAL1* activation events as follows: i) 223 (57%) with *STIL-TAL1* deletion, ii) 58 (15%) by *TAL1* with upstream enhancer indels, iii) 42 (11%) with chromosomal translocations, iv) 28 (7%) with *TAL1* intron enhancer alterations, v) 25 (6%) with *TAL1* downstream enhancer I alterations, and vi) 15 (4%) with *TAL1* downstream enhancer II alterations (**Fig. 1b** and **Extended Data Fig. 1a-b**).

*TAL1* is known to be expressed by normal HSCs and other hematopoietic progenitors but silenced during thymocyte development^24–26^. This was corroborated by *TAL1* expression in single cell RNA-seq (scRNA-seq) from an integrated dataset of both normal bone marrow and normal thymus cells generated by Cordes et al.^26^. Specifically, the highest *TAL1* expression occurred in HSCs, which gradually decreased in the later progenitors: i) MPPs, ii) lymphoid-primed multipotent progenitors (LMPPs), and iii) common lymphoid progenitor (CLP) cells (**Fig. 2a**). By contrast, *TAL1* was not expressed in thymocytes from the normal thymus, and it was previously observed to show expression only at early DN-stages of thymocyte development^24,27^, and interestingly its aberrant activation at later stages is a leukemogenic event in T-ALL^15,24^. Indeed, almost all T-ALLs harboring these somatic variants exhibit high *TAL1* expression (**Fig. 2b**) including the two index cases studied here (SJTALL002048_D (PARSNX): 7.35 TPM, and SJALL015708_D (PAUAZV): 3.97 TPM).

**Fig. 2:**
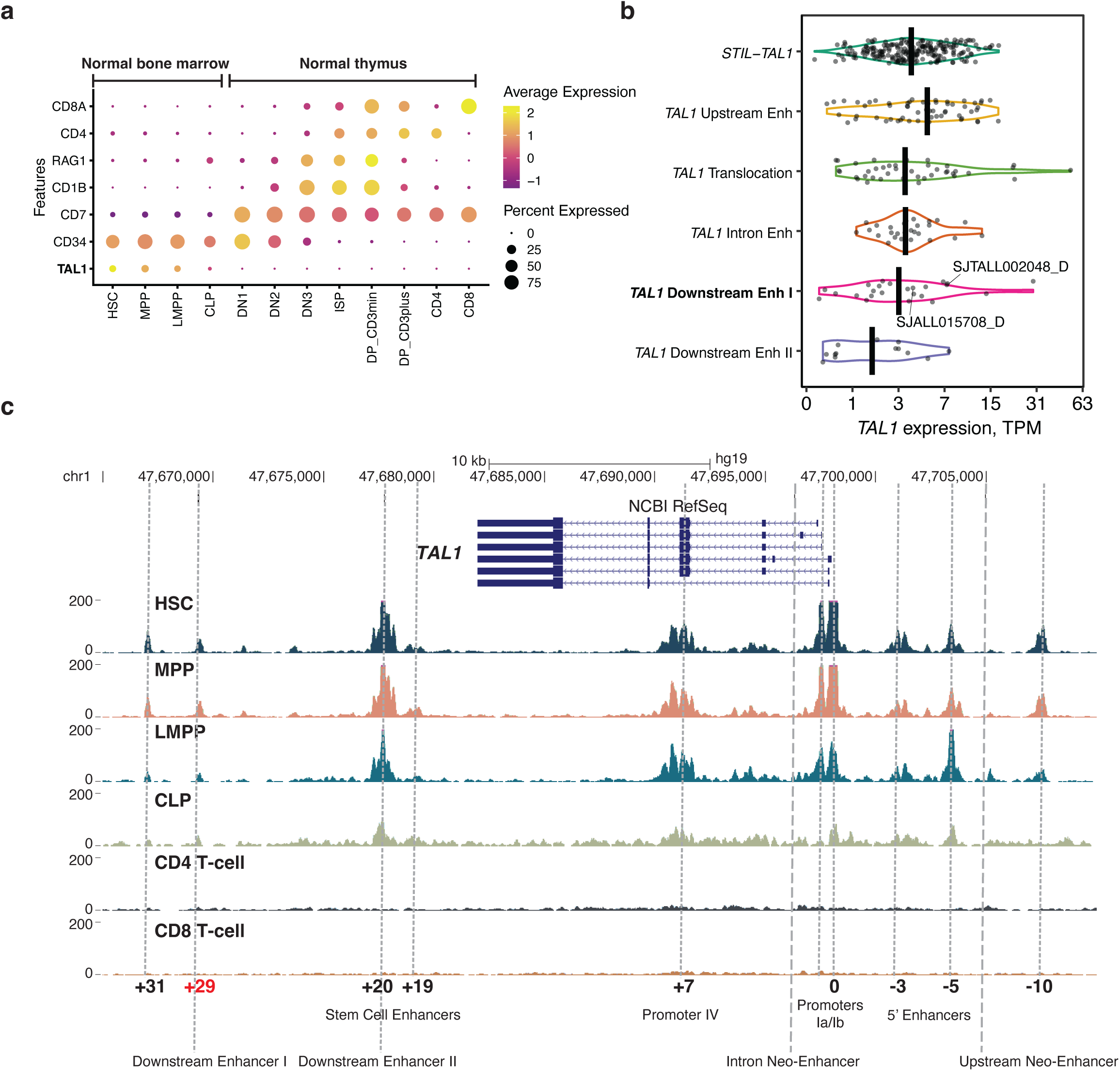
***TAL1* expression and enhancer landscape in T-ALL and normal hematopoiesis. a**, Dotplot showing scaled RNA-seq expression levels of marker genes in normal cells from human bone marrow: i) hematopoietic stem cells (HSCs), ii) multipotent progenitors (MPPs), iii) lympho-myeloid primed progenitors (LMPPs), iv) common lymphoid progenitors (CLPs); and in normal cells from human thymus: i) three stages of CD4-CD8- double negative (DN), ii) immature single-positive (ISP), iii) CD4+CD8+ double-positive (DP) that either express CD3 (CD3+, CD3plus) or not (CD3-, CD3min), and iv) CD4+ or CD8+ single-positive thymocytes. The single cell RNA-seq dataset and cell type annotations were obtained from the study of Cordes et al. 2022^26^. **b**, Violin plot showing *TAL1* RNA-seq expression by T-ALL samples from Polonen et al.^12^ (see Methods). Samples are colored by the type of genomic alterations that mediated *TAL1* activation. Two samples from the index patients from which PDX models were derived in this study are labeled SJTALL002048_D and SJALL015708_D. **c**, Coverage plot showing ATAC-seq data for sorted normal cell populations of HSCs, MPPs, LMPPs, CLPs, CD4 and CD8 T-cells. ATAC-seq data are from Corces et al.^28^, and visualized using the UCSC genome browser (see Methods). ATAC-seq peaks in progenitor cells are denoted by vertical gray dashed lines (small dashes). Three rows at the bottom mark the locations of 1) open chromatin regions defined by presence of ATAC-seq peaks in progenitor cells are labeled by their kbp distance to the promoter 1a based on conventional labelling^10,33^; 2) well-studied enhancer and promoter regions in normal hematopoiesis; and 3) T-ALL enhancers defined by somatic non-coding variants^12^. The two enhancers that do not have open chromatin in progenitor cells are labeled as “Neo-enhancer” and marked with gray dashed lines (larger dashes). On top, the track shows *TAL1* isoforms from NCBI RefSeq.

Mirroring the *TAL1* expression in normal hematopoiesis, ATAC-seq data generated from sorted populations of primary blood cells from healthy individuals^28^ also show multiple peaks in normal hematopoietic progenitor cells where *TAL1* is expressed, which are absent in CD4+ or CD8+ single-positive mature T-cells where *TAL1* is silenced (**Fig. 2c**). In both the HSCs and normal hematopoietic progenitors, ATAC-seq peaks are present at known *TAL1 cis*-regulatory regions, which include i) promoter regions at exons 1 and 4, ii) the upstream −3 enhancer and iii) the downstream +19/+20 stem cell enhancer regions. Accessible open chromatin is also present by this technique in regions with less well-defined enhancer activities, including downstream at +29, +31, as well as upstream at −5 and −10 relative to the promoter. Among the four T-ALL enhancer regions initiated by non-coding variants, the intron 1 and upstream enhancers do not overlap with ATAC-seq peaks in normal progenitor cells, indicating that these sequences do not function as *TAL1* enhancers in normal progenitor cells. By contrast, the T-ALL downstream enhancer II matches the well-characterized +19/20/21 hematopoietic stem cell enhancer, while downstream enhancer I, the region of our interest, overlaps with the +29 ATAC-seq peak present in the normal hematopoietic progenitors.

### Distribution of non-coding variants at *TAL1* downstream enhancer I region

The *TAL1* downstream enhancer I region in T-ALL was previously denoted to have three somatic variant types, which are indels, duplications and inversions^12^. Using the raw WGS data for variant curation, we found that all variants should be reclassified into ITDs and indels (**Extended Data Fig. 1c-d** and **Supplementary Table 1**) which are present in 17 and 8 T-ALLs, respectively (**Fig. 3a-d** and **Extended Data Fig. 1a-b**). Thus, the two T-ALLs selected for this investigation are representative of the two different types of variants in the *TAL1* downstream enhancer I region.

**Fig. 3:**
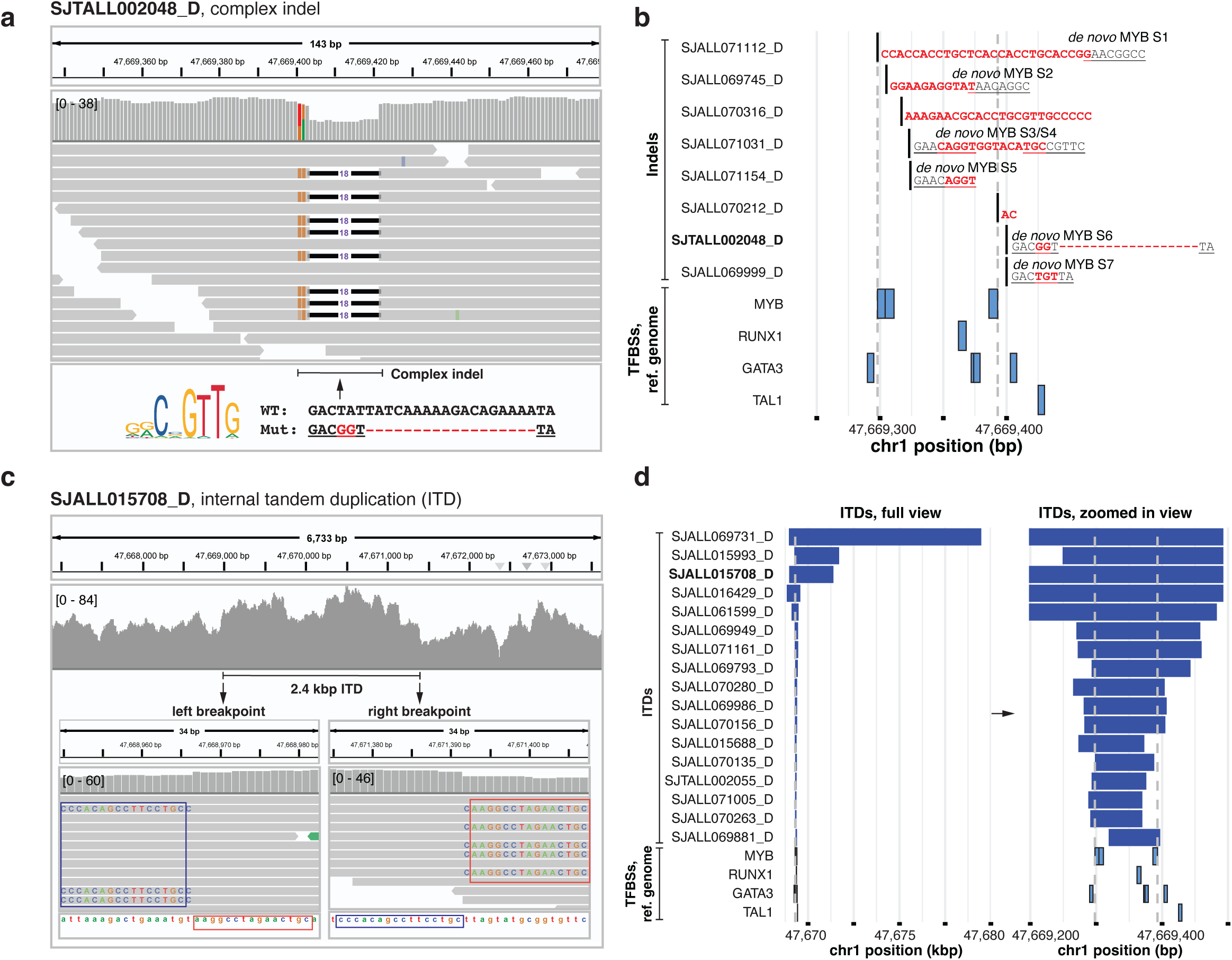
Non-coding somatic variants in T-ALL at the *TAL1* downstream enhancer I region. **a**, WGS reads of a heterozygous complex indel in the index sample SJTALL002048_D displayed using the IGV browser. At the bottom, the MYB binding motif (MA0100.1) from the JASPAR2024 database is shown on the left, and wild type (WT) and mutated sequences are shown on the right with the *de novo* MYB transcription factor binding site (TFBS) underlined. Somatic complex indel alteration is highlighted in red. **b**, Genomic location of somatic indels identified in the Polonen et al. cohort^12^. The coordinates are based on GRCh37 (hg19). Each row represents one sample with nucleotide bases introduced by the somatic indel highlighted in bold red and the reference bases shown in black. *De novo* MYB binding sites introduced by the indel are labeled as MYB S1-7 with the TFBS underlined. The TFBSs for TAL1 CRC members (MYB, RUNX1, GATA3, and TAL1) on the reference human genome are shown at the bottom. The dashed vertical gray lines define a near 100 bp genomic segment where three canonical MYB TFBSs exist. **c**, WGS reads of a heterozygous internal tandem duplication (ITD) in the index sample SJALL015708_D displayed by the IGV browser. On top, increased genomic coverage caused by the 2.4 kbp ITD. At the bottom, left and right ITD breakpoints shown in base-pair resolution marked by blue and red squares highlighting the soft clipped subsequences caused by the ITD breakpoints. **d**, Genomic location of somatic ITDs identified in the Polonen et al. cohort. The left panel is 10 kbp region showing the full range of ITDs while the right panel is a zoomed-in view of 300 bp centered at the minimum overlapping region of ITDs. The two index samples for which PDX models were derived in this study are highlighted in bold in **b** and **d**.

In contrast to the recurrent hotspots for indels and SNVs identified in the other three enhancer regions (**Extended Data Fig. 1b)**, indels within downstream enhancer I region were dispersed over a 103 bp interval (**Fig. 3b**). The ITDs were of variable size (79 - 10,520 bp), with their minimum overlapping region falling into the same interval (**Fig. 3d**). Within the reference human genome, the slightly larger region (141 bp) centered around this interval contains three MYB TFBSs (see Methods and **Supplementary Table 2a**) along with four GATA3, one RUNX1, and one TAL1 TFBSs (**Extended Data Fig. 1f-g**). Interestingly, minor peaks that overlap MYB, GATA3, and RUNX1 TFBSs can be found in ChIP-seq of these transcription factors generated from the Jurkat cell line, despite absence of somatic variants in this enhancer I region (**Extended Data Fig. 1e**)^15^. No H3K27ac super-enhancer peak could be found in the downstream enhancer I region in Jurkat cells, showing that residual binding of MYB and other TFs to the wild type (WT) allele is insufficient for enhancer activation in this region.

Of the two index T-ALLs, SJTALL002048_D harbored a somatically acquired complex indel comprised of double substitution chr1:47,669,401TA>GG and an 18 bp deletion at chr1: 47,669,404 (**Fig. 3a**). This complex indel resulted in creation of a *de novo* MYB TFBS, similar to previously reported recurrent indels at the upstream enhancer region in Jurkat and Molt-3/4 cell lines and patient T-ALL samples^15,29^. Of the eight indels identified in the T-ALL cohort^12^ (**Fig. 3b** and **Extended Data Fig. 1f**), six created *de novo* MYB TFBSs (**Supplementary Table 2b-c**). By contrast, *de novo* MYB binding sites were present only in two out of 17 ITDs (12%, samples SJTALL002055_D and SJALL069881_D, **Supplementary Table 2d-e**) at the ITD junction but absent in the other ITD samples including the index sample SJTALL015708_D which had a 2.4 kbp long ITD (**Fig. 3c** and **Extended Data Fig. 1g**). Among the 17 ITD samples (**Fig. 3d** and see Methods), most (12 out of 17, 71%) had a relatively short length (less than 200 bp), while three had ITDs exceeding 2 kbp.

### Aberrant *TAL1* transcription via downstream enhancer I

Allele-specific expression (ASE) is a robust read-out for transcription activation caused by non-coding *cis*-regulatory variants with heterozygous genotype, including alleles containing somatically introduced alterations, such as indels or ITDs selected for during leukemic transformation^16^. To validate *cis*-activation of *TAL1* expression in the two index T-ALLs, we first evaluated mono-allelic expression of germline single nucleotide polymorphisms (SNPs) in RNA-seq data generated from PDX (Methods) and patient samples. For SJTALL002048, there were six heterozygous expressed SNPs at the *TAL1* locus with sufficient RNA-seq coverage in the PDX sample and five heterozygous SNPs with sufficient coverage in the patient RNA-seq sample. These SNPs were heterozygous in both tumor and matched normal DNA but mono-allelic in tumor RNA-seq of PDX (**Fig. 4a** and **Supplementary Table 3**) and patient sample (**Extended Data Fig. 2a**), confirming ASE. Such analysis could not be performed for SJALL015708 due to lack of heterozygous SNPs at *TAL1* locus for this patient (see Methods).

**Fig. 4:**
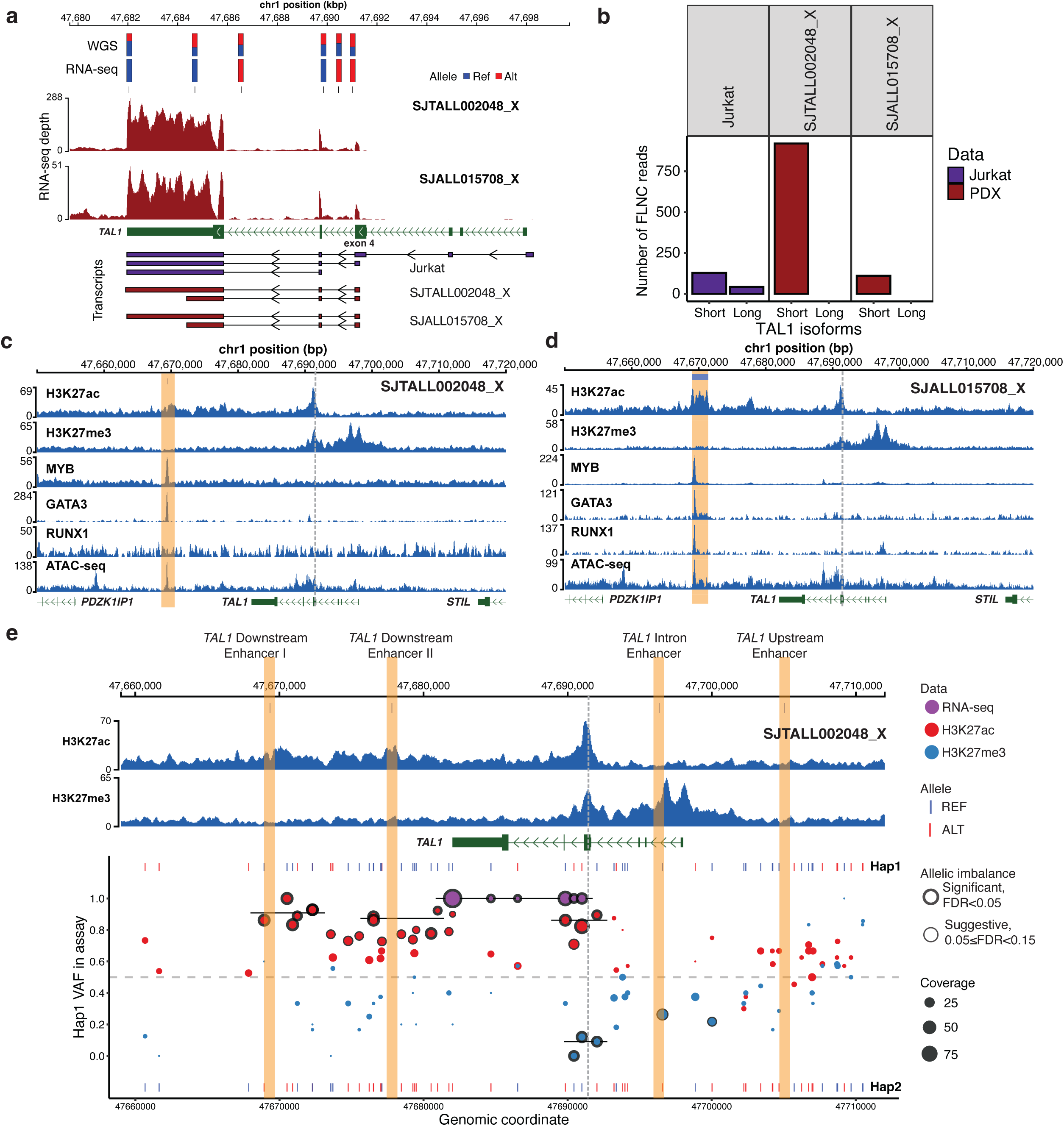
***TAL1* transcription and epigenetic profiles generated from the PDX models. a**, In the middle, *TAL1* RNA-seq coverage from the indel sample SJTALL002048_X (first track) and from the ITD sample SJALL015708_X (second track). Barplots on top show variant allele frequencies (VAFs) of REF (reference, blue) and ALT (alternative, red) alleles of six heterozygous SNPs in SJTALL002048_X which are bi-allelic in tumor DNA (observed in WGS), but mono-allelic in RNA (observed in RNA-seq). At the bottom, plot showing representative transcripts obtained from PacBio Iso-seq for Jurkat cell line and two PDX samples. All transcripts from PacBio Iso-seq are shown in **Extended Data** Fig. 2b. **b**, Comparison of the *TAL1*-long and *TAL1*-short isoforms expressed in the Jurkat cell line with those in the two PDX models samples. Transcription was measured by full-length non-chimeric (FLNC) reads generated from PacBio full-length transcriptome sequencing. A transcript is considered a long isoform unless its transcription start site is located downstream of intron 3, in which case it is annotated as the short isoform. **c-d**, Coverage plots showing ChIP-seq data for the histone modifications H3K27ac and H3K27me3, and for transcription factor (MYB, GATA3 and RUNX1) binding to DNA from the indel sample SJTALL002048_X (**c**) and the ITD sample SJTALL015708_X (**d**). Locations of non-coding mutations in the downstream enhancer I regions are highlighted by orange bands, and the location of *TAL1*-short isoform TSS at exon 4 is denoted with the grey dashed line. The boundaries of ITD are denoted with the blue line at the top in **d**. **e**, Haplotype-phased allelic imbalance in H3K27ac and H3K27me3 ChIP-seq and RNA-seq of the indel sample SJTALL002048_X. The top panel shows the ChIP-seq coverage and the *TAL1* gene structure while the bottom panel shows haplotype-phased allelic imbalance. In the bottom panel, Hap1 and Hap2 are the two haplotypes constructed from germline SNPs of SJTALL002048_D with their phased reference (REF) and alternative (ALT) alleles for heterozygous SNPs shown in blue and red, respectively. Hap1 and Hap2 are arbitrarily designated for the phased variant sites. They are displayed at the top and bottom, respectively, and a horizontal dashed gray line marks the VAF of 0.5 expected for a heterozygous SNP. Each colored dot represents the VAF of a heterozygous SNP in RNA-seq (purple), the active mark H3K27ac (red), and the repressive mark H3K27me3 (blue). The size of the dot corresponds to the data coverage, and significant allelic imbalance is marked with black outline. The significance was evaluated using two-sided binomial test, and the resulting P-values were adjusted with the FDR method. The median Hap1 VAF of a region with consecutive significant SNPs (false discovery rate (FDR) < 0.05) is shown by a horizontal black line. Vertical bands highlight the locations of four T-ALL enhancers.

*TAL1* is known to have two isoforms: *TAL1*-long initiating from the TSS of exon 1, and *TAL1*-short initiating from a TSS embedded within exon 4^30^ (also referred as exon 3 in some studies^31^) of the *TAL1*-long isoform. Interestingly, the RNA-seq of the two PDX samples indicated exclusive expression of the *TAL1*-short mRNA, which included only sequences from exons 4-6 of the long isoform (**Fig. 4a**). To validate this, we performed full-length transcriptome sequencing with PacBio platform of RNA from the two PDX samples (SJTALL002048_X and SJALL015708_X), along with RNA from the Jurkat cell line as a control (Methods). Only *TAL1*-short was detected in the two PDX samples in contrast to the expression of both *TAL1*-short and *TAL1*-long in Jurkat cell line (**Fig. 4b** and **Extended Data Fig. 2b-c**). In both PDX samples, *TAL1* transcription is initiated in close proximity to what was previously referred as the alternative promoter IV or third *TAL1* promoter^30^ (**Extended Data Fig. 2d**), leading to the production of a truncated protein compared to the product of the *TAL1*-long isoform.

### Mechanism of *TAL1* aberrant activation via downstream enhancer I

To better understand the mechanism of *TAL1* activation via downstream enhancer I, we interrogated H3K27ac and H3K27me3 ChIP-seq data generated for the two PDX samples to gain insight into the chromatin modifications accompanying expression of the *TAL1* short isoform (**Fig. 4c-d** and **Extended Data Figs. 3-5**). H3K27ac peaks were present at *TAL1* downstream enhancer I and at the alternative promoter region for the short *TAL1* RNA, which overlaps with exon 4 of the long *TAL1* RNA, in both the SJTALL002048_X and SJALL015708_X samples, but absent in upstream regulatory regions for the long *TAL1* RNA, thereby validating downstream enhancer I activity (**Fig. 4c-d**). Repressive mark H3K27me3 was present in a broad peak over the regions of the upstream enhancer across exon 1 of the long *TAL1* isoform, consistent with repression of expression of the long *TAL1* RNA in both samples (**Fig. 4c-d**). H3K27ac modifications were present over the promoter region of the short *TAL1* RNA in both PDX samples, consistent with high levels of expression of this isoform in leukemias with *cis*-activation of downstream enhancer I, along with evidence for the H3K27me3 modification over this region, likely affecting the reference allele that lacked an activating indel or ITD.

To investigate the involvement of members of CRC in *TAL1* activation^22^, we examined ATAC-seq and ChIP-seq profiles of MYB, GATA3 and RUNX1 generated from the two PDX models (**Fig. 4c-d** and **Extended Data Figs. 3-6**). Globally, co-localization of binding of these CRC TFs was found in both samples (**Extended Data Fig. 7a),** consistent with the binding of the TAL1 CRC members to the allele harboring the activating indel or ITD^15^. Across the region encompassing *TAL1* gene-coding and *cis*-regulatory sequences, the sole prominent peaks of all TFs and ATAC-seq accessible open chromatin occurred at the downstream enhancer I region in both samples (**Fig. 4c-d**), facilitating enhancer activity due to binding of MYB and the other TAL1 CRC transcription factors. This stretch of open chromatin was aligned to the about 100 bp region where the reference human genome contained the canonical TFBSs: three MYB sites, four GATA3 sites, one RUNX1 site, and one TAL1 site (**Extended Data Fig. 5b**). In addition, ATAC-seq accessible open chromatin was evident surrounding the promoter region that drives expression of the short *TAL1* RNA isoform, which corresponds to the location of exon 4 sequences of the long form of the *TAL1* transcript. Co-occupancy of MYB, GATA3 and RUNX1 TFs was found in the ChIP-seq of ITD sample SJALL015708_X, but only MYB and GATA3 peaks were present in the ChIP-seq data of indel sample SJTALL002048_X (**Fig. 4c-d**), indicating that co-occupancy of RUNX1 is not always required for *TAL1* enhancer activity as proposed by a recent study^32^.

We next evaluated the contribution of the mutant allele in TF binding for the ITD sample SALL015708_X. As the ITD junction does not create a new TF binding site and is about 400 bp away from the ATAC-seq region of open chromatin and TFs ChIP-seq peaks, binding specificity to the mutant allele cannot be directly evaluated in the ITD sample SJALL015708_X. By contrast, in indel sample SJTALL002048_X, the indel allele creates a *de novo* consensus MYB binding site at the position where the MYB peak was centered in the ChIP-seq data, and the indel allele was significantly enriched accounting for 77% of the MYB ChIP-seq reads (**Extended Data Figs. 7b-c**). GATA3 binding and the ATAC-seq region of open chromatin were similarly enriched for the indel allele, accounting for 78% and 88% of the reads, respectively (**Extended Data Figs. 7b-c**). The enrichment for the indel allele is consistent with the preferential TF binding results for the mutant allele at the upstream enhancer region in Jurkat cells^15^, where the indel-containing reads accounted for 96% and 92% reads at regions enriched for MYB and GATA3 binding, respectively (**Extended Data Figs. 7d-e**). The enrichment for the mutant allele in the indel sample indicates that the enhancer activity was driven by the introduction of the new MYB binding site in the complex indel despite the presence of a canonical MYB binding site in close proximity (**Extended Data Fig. 1f**). Residual MYB and GATA3 bindings at the enhancer I region, initially observed in Jurkat cells (**Extended Data Fig. 1e**), were replicated by the 22-23% MYB and GATA3 bindings contributed by the WT allele in the indel sample SJTALL002048_X (**Extended Data Fig. 7e**).

### Haplotype specificity of H3K27ac and H3K27me3 binding across the *TAL1* locus

For the indel sample SJTALL002048_X, the enrichment for TF binding to the mutant allele indicates that the mutant allele drove the enhancer activity. Indeed, H3K27ac histone modification at the indel site showed an enrichment for the mutant allele (70%), but its significance could not be assessed due to insufficient coverage at the indel site. Furthermore, as the downstream enhancer I region is separated from the *TAL1* promoter IV by 22 kbp, a direct connection between the enhancer and promoter activities could not be established. Therefore, we explored the potential to make such a connection through haplotype phasing (Methods). This analysis is feasible only for the indel sample SJTALL002048_X which had 60 heterozygous SNPs, but not for the ITD sample, which only had two heterozygous SNPs across the region (**Supplementary Table 3**). By arbitrarily selecting one of the two constructed haplotypes as the variant allele (i.e. Hap1, **Fig. 4e**), we examined allelic imbalance in RNA-seq as well as in ChIP-seq assays for the active mark H3K27ac and the repressive mark H3K27me3 (**Fig. 4e**).

The most prominent allelic imbalance in the H3K27ac modification (red dots, **Fig. 4e**) was found at a 5 kbp region encompassing downstream enhancer I which was enriched for the Hap1 allele with five consecutive SNPs consistently exhibiting significant enrichment (**Fig. 4e**). Thus, the somatically acquired complex indel in this region, also enriched in H3K27ac, most likely occurred on the Hap1 allele. Significant enrichment for Hap1 in H3K27ac was also found at downstream enhancer II as well as at exon 4 where *TAL1* transcription was initiated from the alternative promoter IV. Notably, the repressive mark H3K27me3 (blue dots, **Fig. 4e**) was significantly enriched for the other haplotype, Hap2. This shows that the co-existence of the active H3K27ac peak and repressive H3K27me3 peak at exon 4, although paradoxical at first glance, actually represented the different chromatin states of the two haplotypes in this T-ALL. Mono-allelic expression detected by RNA-seq (purple dots, **Fig. 4e**) also showed a perfect match to Hap1, consistent with the active state of Hap1 and repressed state of Hap2 based on the allelic imbalance in histone marks.

The presence of an active downstream enhancer and an active promoter at exon 4 on the same haplotype suggests that promoter activity may be driven by enhancer-promoter contact. The exon 4 promoter was previously found to contact downstream enhancer II in the Jurkat cells^33^. Capture Hi-C data generated from progenitor CD34+ cells^34^ also showed contact between the downstream enhancer I region and the *TAL1* exon 4 promoter, but not the exon 1 promoter (**Extended Data Fig. 8a**). To evaluate whether such interaction exists in the T-ALLs immortalized as two PDXs under study, we generated Hi-C data from these two PDX samples for comparison with the two cell line controls, i) Jurkat cells (*TAL1*-positive, activated via upstream enhancer alteration) and ii) DND41 cells (*TAL1*-negative). At the *TAL1* locus, higher contact density was observed in the two PDX samples compared to the DND41 cell line, and the Jurkat cell line also showed similarly high density of DNA contacts, which is consistent with its active *TAL1* state (**Extended Data Fig. 8b**). Furthermore, there was an enriched coverage corresponding to the contacting regions overlapping exon 4 and the downstream enhancer region in our two PDX samples, which was absent in the DND41 cell line (**Extended Data Fig. 8b**). Because of the low resolution of Hi-C, downstream enhancer I and II regions at the *TAL1* locus cannot be distinguished. These results support the hypothesis that transcription of the *TAL1*-short isoform is initiated by DNA looping of the reactivated downstream enhancer I, such that it contacts the exon 4 promoter in our two T-ALL PDX samples.

### Prediction of *TAL1* enhancer variant effect by AlphaGenome

In a recent published study, AlphaGenome^23^, a sequence-to-function AI model, was used to predict functional impact of published *TAL1* non-coding variants in upstream enhancer^15^, intron enhancer^16^ and downstream enhancer II^17^, but not those in downstream enhancer I. Here, we performed AlphaGenome prediction on all four *TAL1* enhancer regions to evaluate whether its performance on downstream enhancer I variants was comparable to those on the variants from other three enhancer regions.

We first compared AlphaGenome’s predicted *TAL1* expression scores with the observed RNA-seq expression level in 122 T-ALLs with *TAL1* activation driven by non-coding variants in these four enhancer regions analyzed in this study (see Methods). Jurkat cell line was also analyzed to serve as the positive control, which, as expected, showed a high concordance between the predicted and the observed tracks of RNA expression and H3K27ac ChIP-seq (**Extended Data Fig. 10)**. For patient T-ALL samples, those harboring variants in upstream enhancer and downstream enhancer II had high predicted *TAL1* RNA expression scores, consistent with observed expression level by RNA-seq (**Fig. 5a-b**). T-ALLs activated by the recurrent SNV in intron enhancer only had a single predicted score (the variant recurrence is indicated by the size of the dot in **Fig. 5a**) despite varying observed RNA-seq expression and therefore were not included in this comparison. T-ALLs with *TAL1* activated by variants in downstream enhancer I, our region of interest, had significantly lower predicted scores compared to those with variants in upstream enhancer and downstream enhancer II (Wilcoxon rank-sum two-sided test P = 3.32×10^-7^). The same trend persisted when considering only the indel variants, as they corresponded more closely to the SNV/indel variants of 1-20 bp length used in the AlphaGenome’s distillation training process (**Extended Data Fig. 10a)**. By contrast, observed *TAL1* RNA-seq expression values in T-ALLs with *TAL1* activated by enhancer I region variants were comparable to those activated by the other two enhancers’ variants (**Fig. 5b** and **Extended Data Fig. 10b**, Wilcoxon rank-sum two-sided test P = 0.35). Thus, the predicted expression scores for variants in enhancer I region did not match the observed RNA-seq data. Of note, similar to T-ALLs with *TAL1* activated by the intron enhancer, recurrent variants in these three enhancer regions also exhibited variable RNA-seq expression level despite having the same prediction scores by AlphaGenome, suggesting that additional factors may contribute to *TAL1* expression variation in T-ALL.

**Fig. 5:**
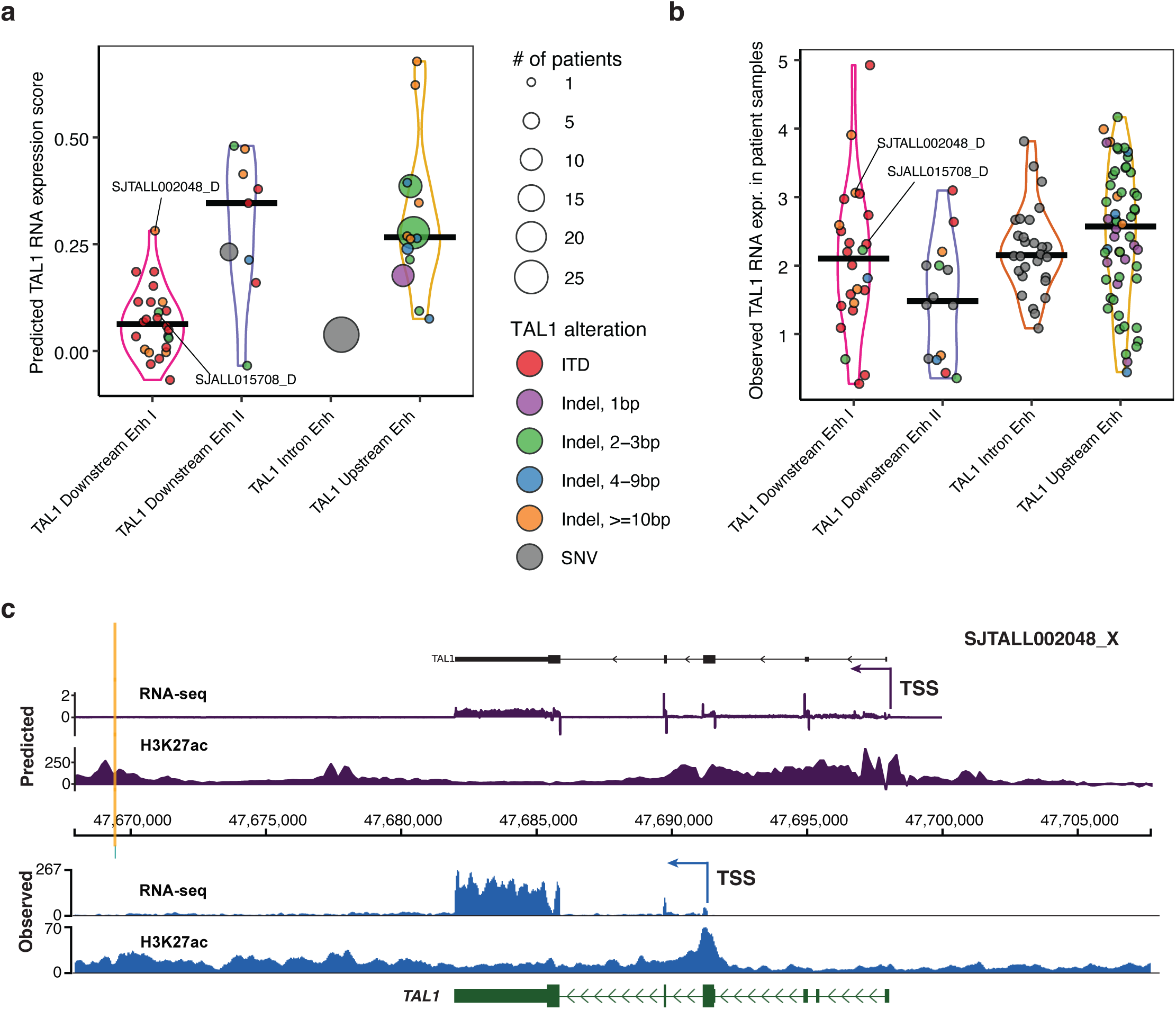
Predictions of *TAL1* variants impact with AlphaGenome. **a**, Violin pot showing distribution of the *TAL1* RNA expression scores predicted by AlphaGenome for the SNV, indel, and ITD variants that affect one of the four enhancer regions mediating *TAL1* expression in 122 T-ALL patient samples. Dot size corresponds to variant recurrence, i.e. the number of T-ALLs with the same variant. **b**, Violin plot showing distribution of *TAL1* RNA-seq expression (log_2_(TPM+1) values) in 122 T-ALLs with variants affecting *TAL1* expression that are shown in panel **a**. **c**, Predicted and observed coverage tracks for *TAL1* RNA-seq and H3K27ac ChIP-seq in SJTALL002048_X sample with indel affecting *TAL1* downstream enhancer I. On top, coverage tracks show predicted by AlphaGenome *TAL1* RNA-seq expression and H3K27ac ChIP-seq scores (difference between ALT and REF sequences). At the bottom, coverage tracks show *TAL1* expression and H3K27ac ChIP-seq signal from the experimental data. Arrows mark the *TAL1* TSS at exon 1, and alternative *TAL1* TSS at exon 4.

AlphaGenome also provides the predictions of variant effects on RNA-seq and ChIP-seq tracks at a single-base-pair resolution^23^. For Jurkat cell line, its predicted RNA-seq and H3K27ac ChIP-seq tracks showed high concordance with the real data from the experimental assays with high coverage of H3K27ac and RNA-seq at exons 1 and 4 (**Extended Data Fig. 10c**), supporting transcription initiation at these two sites leading to expression of both the long and the short *TAL1* isoforms. However, such concordance was absent when comparing the predicted tracks with observed data generated from the PDX samples of our two index cases with *TAL1* downstream enhancer I variants (**Fig. 5c** and **Extended Data Fig. 10d**). Transcription of the short isoform initiated within exon 4, which was evident in experimental RNA-seq and H3K27ac ChIP-seq generated from PDX models, was absent in the predicted tracks for both cases. While predicted RNA expression score for the indel case (SJTALL002048), which was the highest among all variants in downstream enhancer I region (ranked 3^rd^ in the observed RNA-seq data, **Fig. 5a-b**), did support *TAL1* activation, the prediction of ITD sample (SJALL015708) even had a low positive score of 0.08, showed changing pattern of activation/deactivation across *TAL1* gene body (**Extended Data Fig. 10d**).

### Molecular features of T-ALLs with *TAL1* activated by downstream enhancer I

Because the downstream enhancer region is active in hematopoietic progenitor cells, we first examined whether samples with *TAL1* expression activated via downstream enhancer I show expression patterns characteristic to cells undergoing normal T-cell development stages in the thymus. *TAL1* T-ALLs were reported to have two distinct subtypes: i) *TAL1* double-positive-like (DP-like) corresponding to the DP stage of normal thymocyte development; and ii) *TAL1* alpha/beta-like corresponding to more mature single positive alpha/beta expressing normal thymocytes, which are characterized by the expression of *TRAC* and the presence of TCR alpha/beta DNA rearrangements^12^. These two subtypes have differentially expressed immune markers, i) *CD1E*, *CD1B*, *CD4* and *CD8*, which are highly expressed in *TAL1*-expressing DP-like T-ALL subtype, and ii) *TRAC*, *PTPRS* and *IGFBP5*, which are highly expressed in *TAL1* alpha/beta-like T-ALL subtype (Methods, **Extended Data Fig. 9a-b** and **Supplementary Table 4A)**. Here we examined the appearance of these two *TAL1* T-ALL subtypes within the six T-ALL groups based on the mechanism of *TAL1* activation (**Fig. 2b**). Interestingly 22 out of 25 (88%) *TAL1* T-ALLs from the downstream enhancer I group were assigned a *TAL1* DP-like subtype (Fisher’s exact test FDR = 8.94×10^-5^, **Fig. 6a** and **Extended Data Fig. 9c**), indicating an earlier developmental stage. The two *TAL1* downstream enhancer I samples studied here were both of *TAL1* DP-like subtype, and they exhibited a characteristic pattern of high expression of *TAL1* DP-like subtype markers, both in patient tumor samples and all T-ALL samples from their respective PDXs (**Extended Data Fig. 9d**). We also evaluated the expression of these markers by the Jurkat cells. Interestingly, the Jurkat cell line exhibited expression of markers from both subtypes (**Extended Data Fig. 9d**). As expected, members of *TAL1* CRC, such as *MYB*, *TCF12*, *RUNX1*, *CREBBP*, and *GATA3*^22^ are all highly expressed in patient T-ALL cells, PDXs and Jurkat cell line samples.

**Fig. 6:**
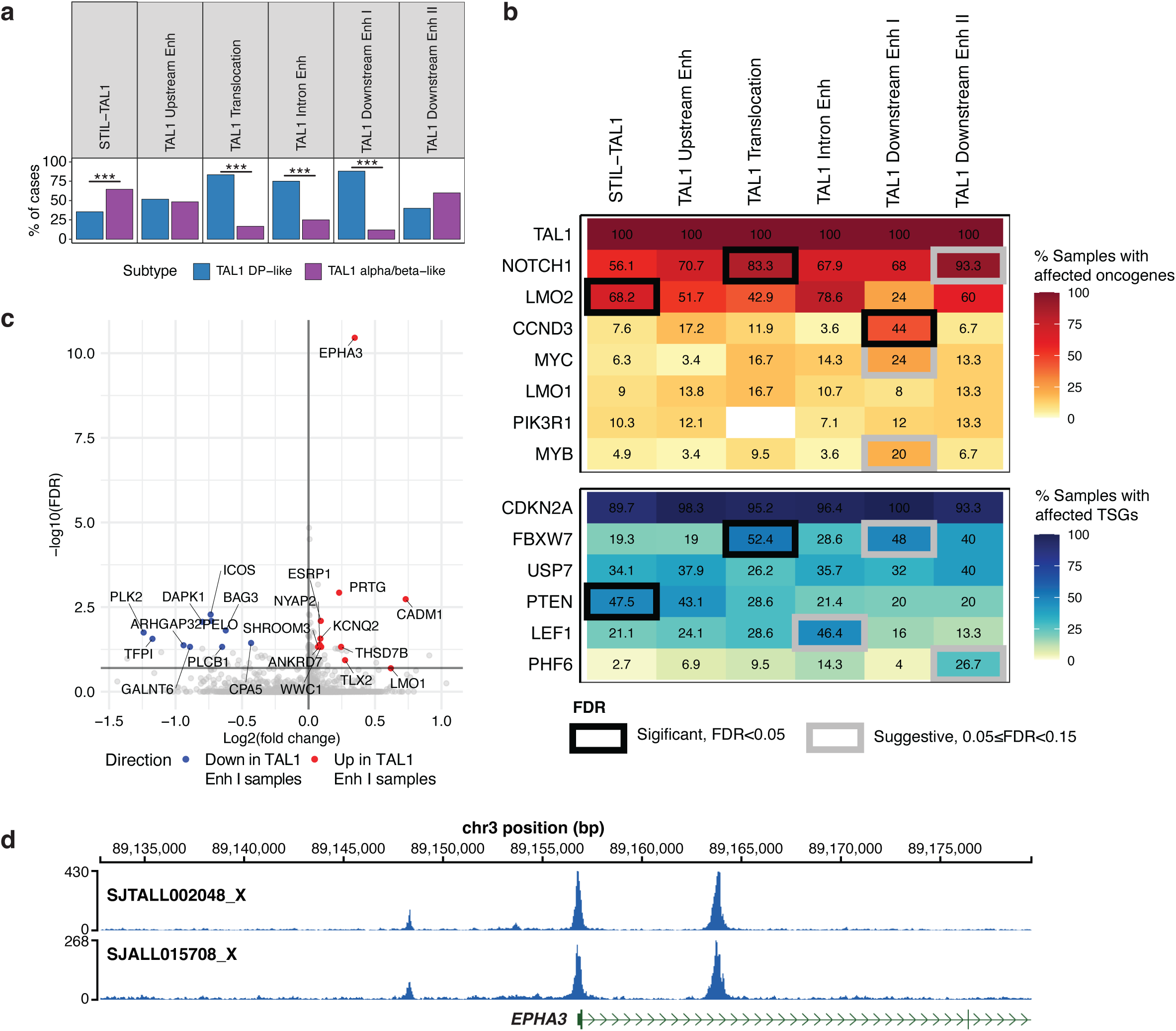
Molecular features of the *TAL1* downstream enhancer I group. **a**, Distribution of DP-like vs alpha/beta-like subtypes across the six groups of samples with *TAL1 cis*-activating alterations. Significant enrichments of one subtype with FDR adjusted Fisher’s exact test P-values < 0.01 are labeled with ***. The *TAL1* downstream enhancer I group is most enriched for DP-like subtype. **b**, Heatmaps showing mutation frequency of T-ALL oncogenes (top heatmap) and tumor suppressor genes (TSGs, bottom heatmap) in the six *TAL1* groups. The black and gray outlines indicate significantly (FDR < 0.05) and suggestively (0.05 σ; FDR < 0.15) enriched gene mutations in a *TAL1* groups, respectively. The significance was evaluated using Fisher’s exact test, and the resulting P-values were adjusted with FDR method. **c**, Volcano plot showing top DEGs between the *TAL1* downstream enhancer I group and the other *TAL1* groups. *TAL1* subtype was used as a covariate in the DEG analysis (see Methods). Top up/down-regulated DEGs are highlighted. Y-axis reflects the −log_10_ of the FDR adjusted P-values from linear model results. Horizontal line corresponds to −log_10_(FDR) = 0.2. **d**, ATAC-seq peaks showing accessibility for promoter region of *EPHA3* in PDX samples SJTALL002048_X and SJTALL015708_X.

Given the enrichment for an earlier DP thymocyte development stage in *TAL1* T-ALLs of the downstream enhancer I group, we then compared their collaborating genomic drivers with the other five groups of *TAL1* T-ALLs (**Fig. 6b**). This group had significant enrichment (Fisher’s exact test FDR < 0.05) for *CCND3*, and marginal enrichment (Fisher’s exact test 0.05 σ; FDR < 0.15) for *FBXW7*, *MYC*, and *MYB*. *CCND3* is a well-known driver affecting cancer cells proliferation, and it is known to be regulated by TAL1 in T-ALL^35^. *FBXW7*, a tumor suppressor protein product of which targets the degradation of oncoproteins, including MYC and NOTCH1^36^, was reported to be repressed by TAL1 via miR-223 in T-ALL^27^. MYB is one of the major TAL1 CRC proteins^22^ and is a critical regulator of thymocyte development^37^. Thus, the *TAL1* T-ALL downstream enhancer I group has enriched representation of mutations or dysregulations of collaborating drivers involved in the *TAL1* gene regulatory network.

Next, to understand transcriptional changes associated with the downstream enhancer I group, we performed differentially expressed gene (DEG) analysis between the samples with *TAL1* activated via downstream enhancer I and samples with other types of *TAL1* activation (**Supplementary Table 4b**). For this analysis, we used *TAL1* subtype (i.e., DP-like or alpha/beta-like) as a covariate, to ensure the DEGs were not driven by the *TAL1* subtype frequency among the groups. We observed 73 DEGs with up-regulated expression and 10 DEGs with down-regulated expression (linear model FDR < 0.05) in the downstream enhancer I group (**Fig. 6c-d** and **Extended Data Fig. 9e**). *EPHA3* was the upregulated gene most significantly associated with the downstream enhancer mechanism regulating *TAL1* expression (linear model FDR = 3.5×10^-11^; **Fig. 6c**). *EPHA3* encodes a receptor protein tyrosine kinase in the ephrin receptor family. Signaling through this receptor in a mouse bone marrow transplantation model was reported to affect normal hematopoietic stem cell adhesion through an integrin-mediated mechanism^38^. The *EPHA3* receptor is expressed by multiple different hematopoietic malignancies including T-ALL, B-ALL and multiple myeloma cells, including by leukemia stem cells^39^.

## Discussion

*TAL1* is a class II bHLH transcription factor that forms heterodimers with E2A or HEB, its class I bHLH partners, to drive gene expression programs that regulate the formation of most hematopoietic cell lineages^8,40,41^. Notably, *TAL1* is essential for establishing the blood program during embryogenesis and for controlling adult hematopoietic stem cell quiescence and survival, as well as for the development of normal erythroid precursors and megakaryocytes. E2A and HEB heterodimers are especially important for the development and differentiation of normal B- and T-lymphocytes and prominently regulate the induction of TCR gene rearrangements in thymocytes, in this case without the involvement or expression of *TAL1*^42^. In fact, the aberrant expression of high levels of *TAL1* in developing thymocytes forms the basis for one of the most prominent genetic subtypes of T-ALL.

Investigators have uncovered at least six complex *cis*-regulatory genetic alterations that are selected for because they lead to the expression of high levels of *TAL1* in developing thymocytes and contribute to the malignant transformation of T-ALL (**Fig. 1a-b**)^11,12,15,17,33^. These mechanisms include chromosomal translocations that hijack active enhancers from the TCR genes into proximity with *TAL1* to drive aberrant expression of *TAL1* and the upstream deletion that places *TAL1* under the control of active enhancers that normally control expression of the *STIL* gene. Also involved are non-coding somatic variants that generate active enhancers within intron 1, as well as in non-coding regions upstream and downstream of *TAL1* coding sequences. These non-coding variants have been shown to generate neo-enhancers that lead to aberrant activation of *TAL1*.

Our experimental studies here have focused on *cis*-activating non-coding variants that have been selected in a region referred to as downstream enhancer I, mediating high levels of *TAL1* expression, that have previously been less studied than the other types of non-coding variants. Using ATAC-seq results published for human normal hematopoietic cell types, we found that non-coding variants known to occur in the two *TAL1* T-ALL downstream enhancer regions are located in regions that are accessible in subsets of normal hematopoietic progenitor cells but not in normal mature T-cells (**Fig. 2c**). In fact, one of these two regions, denoted downstream enhancer II and previously considered to form a neo-enhancer in T-ALL^17^, actually matched the well-studied *TAL1* stem cell enhancer +/19/20/21 active in embryonic and adult hematopoietic progenitors and stem cells^8,43^. This indicates that creation of a neo-enhancer may not always be an underlying mechanism.

Here we propose a new model for aberrant *TAL1* activation in T-ALL, which involves reactivation of a downstream enhancer that is active in HSC and progenitor cells but silenced in T-cells during normal hematopoiesis (**Fig. 7**). Such reactivation in T-ALL is achieved by somatically acquiring variants in the form of indels or ITDs in the same non-coding region where binding sites for TFs of the TAL1 CRC are already present (**Fig. 7**). Studies in normal samples of the relevant hematopoietic lineages can be relied upon for normal characteristics of these sequences in various types of normal hematopoietic progenitors, and these characteristics then compared to the contrasting allelic imbalance patterns of H3K27me3 and H3K27ac binding in T-ALLs with expression of the short *TAL1* isoform initiated from a promoter within exon 4 driven by downstream enhancer I. The indel sample SJTALL002048_X provides support for the hypothesis that this enhancer region, which is normally active in hematopoietic stem cells and early lymphoid progenitors, becomes aberrantly open and active in developing thymocytes due to the presence of the indel sequences. Specifically, enrichment for the repressive mark H3K27me3 on the wild-type haplotype at exon 4 is a footprint of a prior silencing event, while its replacement by the active mark H3K27ac on the mutant haplotype reflects reactivation driven by the non-coding variant (**Fig. 4e**). This type of non-coding variant driven reactivation of enhancer activity is not limited only to the *TAL1* locus or T-ALL or to somatic mutations. For example, prior studies have shown that germline SNPs can deactivate silencers leading to oncogene expression^44,45^, and in B-cell lymphoma 13% of B-cell super-silencers, i.e. regions with significant Polycomb driven H3K27me3 modifications, can be converted to super-enhancers during disease development^46^. Interestingly, *TAL1* was also denoted as a super-silencer locus in the GM12878 cell line derived from B-lymphocytes, a cell type in which *TAL1* is silenced.

**Fig. 7:**
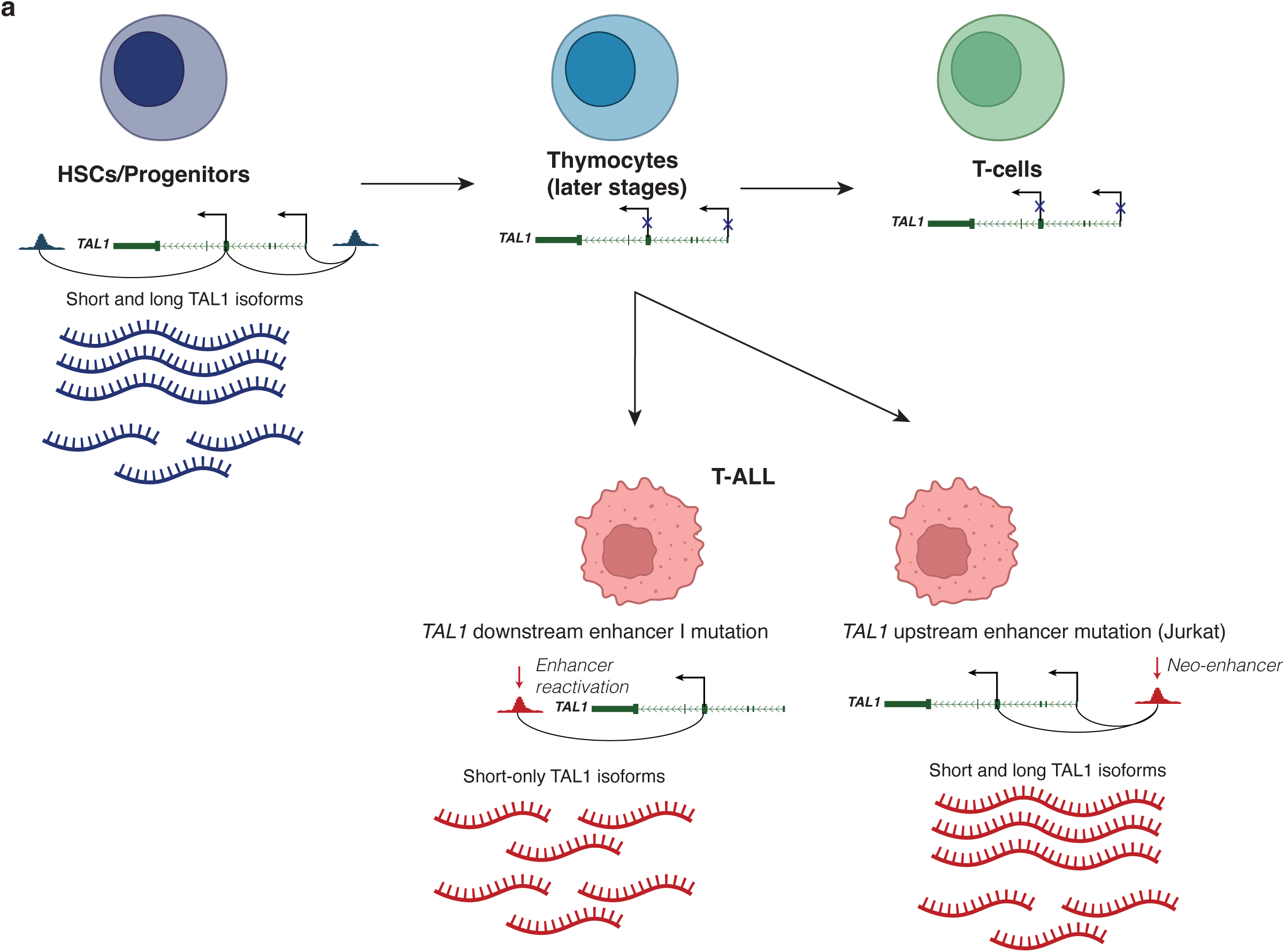
Schematic of *TAL1* enhancer reactivation vs neo-enhancer models. **a**, Top row, *TAL1* regulation in normal hematopoiesis. In normal HSCs and progenitor cells, an active stem cell enhancer located 3’ of the transcribed sequences and upstream enhancer loop to contact the *TAL1* TSS and promoter located at the 5’ origin of exon 4 to enhance the expression of the *TAL1*-short isoform. The *TAL1*-long isoform is also expressed from the TSS at the upstream end of exon 1, which is contacting through looping with the upstream enhancer. In single-positive thymocytes and maturing T-cells before and after exiting the thymus both *TAL1* isoforms are silenced. Bottom row, *TAL1 cis*-activation in T-ALL driven by somatic non-coding variants. On the left, reactivation of the stem cell enhancer by somatic variants at the downstream enhancer I region, leading to expression of the *TAL1*-short isoform, as examined in detail in this manuscript. On the right, an upstream neo-enhancer is created in Jurkat cells and in series of primary T-ALLs^15^, leading to co-expression of both the *TAL1*-short and *TAL1*-long isoforms. Created with BioRender.com.

By curating the data assembled by Polonen et al.^12^, *TAL1* downstream enhancer I was found to be recurrently mutated in 6% of *TAL1* T-ALLs with two types of alterations: ITDs and indels (**Extended Data Fig. 1a-b**). Moreover, while the other three enhancers associated with *TAL1* are activated by recurrent hotspot mutations leading to creation of *de novo* TF binding sites, indels in downstream enhancer I are dispersed across the about 100 bp region where three canonical MYB TFBSs co-exist with canonical GATA3, TAL1, and RUNX1 binding sites in the reference genome. This may render creation of a novel MYB binding site non-obligatory for enhancer activation — because residual MYB binding in this region can be found in Jurkat cells, which do not have any somatic mutations (**Extended Data Fig. 1e**) as well as from the WT allelic fraction of ChIP-seq assays in the indel sample (**Fig. 4e** and **Extended Data Fig. 7e**). Rather, chromatin alterations associated with these variants may alter the accessibility and affinity of binding of these TFs to existing sites, resulting in orders of magnitude greater TF binding levels as well as the formation of an active enhancer. In addition, change of chromatin structure, which could occur due to ITD or indels that do not introduce MYB binding sites, may be crucial for making those binding sites accessible for TFs in the mutated haplotype. Activation of a *TAL1* enhancer requires participation of TAL1 CRC TFs, and the presence of their canonical TFBSs makes it possible for co-occupancy of MYB, GATA3 and RUNX1 at the enhancer region. Interestingly, contact between exon 4 and enhancer I region has also been detected in CD34+ stem cells (**Extended Data Fig. 8A**). These data suggest that the enhancer activity of the *TAL1* downstream region in normal progenitor cells may render it a preferred target for aberrant *TAL1 cis*-activation in T-ALL, because the region already contains the required elements for *TAL1* activation within the reference human genome, i.e. CRC TFBSs and structural features conducive to enhancer-promoter contact.

In a recent study that reports AlphaGenome^23^, a sequence-to-function deep learning AI model that outperforms nearly all other state-of-the-art such models, non-coding variants in the three published *TAL1* enhancer regions (i.e. upstream enhancer, intron enhancer and downstream enhancer II) were used to demonstrate AlphaGenome’s prediction accuracy. Here, we show that unlike the published enhancer regions, *TAL1* RNA expression driven by non-coding variants in the downstream enhancer I region in our study is not well-predicted by AlphaGenome (**Fig. 5a-b**). Furthermore, the predicted tracks of RNA-seq and H3K27ac ChIP-seq did not match the real data generated from PDX models (**Fig. 5c** and **Extended Data Fig. 10D**) as they did not support the transcription of the *TAL1* short isoform observed in real data, even for the indel case which had the highest predicted RNA expression score among the samples with variants in the downstream enhancer I region. This could be in part due to the lack of the representation of the short isoform in GENCODE^47^, the reference database for gene and transcript annotations used for AlphaGenome’s model evaluation. Aberrant activation of oncogenic isoforms not represented in GENCODE is not limited to *TAL1*, as we had previously discovered that two other oncogenes, *FOXR2* and *ERG*, were transcribed from alterative transcription initiation sites in pediatric solid tumor^48^ and leukemia^49^, respectively. These results demonstrate the importance of generating functional data sets, such as RNA-seq and ChIP-seq, in patient samples even for a well-studied gene such as *TAL1*. Incorporation of such data, including those generated by full-length transcriptome sequencing, during the training of AI models such as AlphaGenome can potentially improve the accuracy of predicting RNA transcription, particularly in cases where alternative initiation sites are being utilized.

Our study highlights the importance of non-coding alterations in the activation of expression of the *TAL1* oncogene in T-ALL, revealing a new *cis*-regulatory mechanism leading to reactivation of a lineage-specific enhancer. By integrating publicly available epigenetic data generated from normal hematopoietic cell types with those we have generated here from *TAL1*-transformed T-ALL cells, we show that enhancers with normally restricted activity in specific lineages can be reactivated by genomic alterations. If an enhancer region that is quiescent in normal cells can be activated by somatic variant, the oncogenic configuration will be selected because it will lead to the clonal outgrowth of leukemic cells when it occurs in susceptible progenitors. The comparison of the normal lineage-specific enhancer landscape of hematopoietic stem and progenitor cells with the aberrant enhancer activity observed in leukemic cells represents a powerful approach to uncover mechanisms of transformation in human cancers. The approach we used here can be applied in future studies to derive mechanistic insights into the role of non-coding alterations in malignant transformation while the data that we generated may help improve the predictive ability of AI models such as AlphaGenome.

## Methods

### Specimen data

Tumor samples were collected from two patients diagnosed with T-ALL enrolled on the Children’s Oncology Group (COG) clinical trial, AALL0434 (NCT00408005). Both subjects also enrolled on a companion biology study for biobanking and risk stratification, AALL03B1 (NCT00482352). Genomic studies performed for this work were approved by COG ALL Biology Disease Committee and NCI Cancer Evaluation and Therapeutic Program (CTEP) under the protocol of COG ALL BIOLOGY PROTOCOL AALL17B1-Q. Written informed consent/assent was obtained from the patients and/or their legally authorized representative in accordance with the Declaration of Helsinki.

## Experimental methods

### Cell lines (Jurkat and DND41)

Human T-cell Leukemia cell lines, Jurkat and DND41, were purchased from the American Type Culture Collection (ATCC, Cat# TIB-152) and the Lebniz Institute – German Collection of Microorganisms and Cell Cultures (DSMZ, Cat# ACC 525), respectively. Both cell lines were selected for their relevance in TAL1 activation in pediatric T-ALL.

Jurkat cells were cultured in Gibco RPMI-1640 Medium (Fisher Scientific, Cat# 11875093) supplemented with 10% GenClone Fetal Bovine Serum (FBS) (GeneseeScientific, Cat# 25-514). DND-41 cells were cultured in Gibco RPMI-1640 Medium (Fisher Scientific, Cat# 11875093) supplemented with 10% GenClone heat inactivated Fetal Bovine Serum (h.i.FBS, GeneseeScientific, Cat# 25-514H). Cells were maintained at 37°C in a humidified atmosphere with 5% CO2. The media was replaced every 2-3 days.

Both cell lines were periodically authenticated using short tandem repeat (STR) profiling to ensure they were not misidentified or contaminated with other cell lines. Both cell lines were mycoplasma negative during the culture in this project.

### PDX models

PDX models SJTALL002048_X (CPDM_0075X; PARSNX) and SJALL015708_X (CPDM_0074X; PAUAZV) were generated at the Center for Patient Derived Models at Dana-Farber Cancer Institute, directed by Dr. Keith L. Ligon, using 6-8-week-old NSG mice (NOD.Cg-Prkdc^scid^ Il2rg^tm1Wjl^/SzJ, JAX: #005557). Patient cells were harvested and resuspended in sterile PBS to allow for injection of up to 200 uL per mouse intravenously in the lateral tail vein for p0 PDX generation. Animals were euthanized via CO2 asphyxiation, cervical dislocation, or other Institutional Animal Care and Use Committee (IACUC) approved method. Tumor cells were harvested from the spleen by pulverizing the spleen in RPMI, filtering through 70 micrometer mesh to remove cell clumps and connective tissue, followed by erythrocytes removal using RBC lysis buffer (Qiagen #158902). Cells were washed and viably cryopreserved in FBS with 10% DMSO. Tumor cells were subsequently serially passaged in NSG mice for further expansion and banking.

TALL PDX cells were expanded in the Genomics Lab in the department of Computational Biology (CBGL) in St Jude Children’s Research Hospital (SJCRH). Briefly, NSG mice were bred in the animal research center in SJCRH, each 8-week-old female NSG mouse was injected with ∼10 M of PDX cells through tail veins. Animals were phlebotomized no later than 2-3 weeks after injection and assessed for the presence of circulating disease by peripheral blood flow cytometry. A cell surface marker panel was used in the flow cytometry analysis to detect the tumor cells in the circulation (hCD45-APC-Cy7, hCD19-BV605, hCD2APC, hCD7-PE, mTer119-PerCP-Cy55, mCD45-BUV396, DAPI). Repeat peripheral blood flow cytometry at regular intervals, bi-weekly before detecting tumor cells and weekly after detecting the tumor cell in the peripheral blood, until mice reach the level of the endpoint of harvesting PDX cells, 40∼50 % tumor cells in the peripheral blood. To harvest PDX cells, mice were euthanized at the designated endpoint, splenocytes were harvested and analyzed by flow cytometry to confirm the tumor purity and tumor cell viability. To bank the PDX cells, erythrocytes were eliminated using RBC lysis buffer (Qiagen #158902), and cells were viably frozen in FBS with 10% DMSO.

### RNA-seq data generation

Cell line RNA-seq data was generated using 1 µg of total RNA with the TruSeq Stranded Total RNA Kit (Illumina) as instructed by the vendor protocol using 12 cycles of PCR for library amplification. Sequencing data was generated on an Illumina HiSeq 4000 using the paired end 75 cycle protocol.

PDX RNA-seq data were generated using the KAPA RNA HyperPrep Kit with RiboErase (HMR) (Roche) according to manufacturer’s instructions using 120 ng total RNA input, with the initial incubation at 94 degrees for 4 min, and 12 cycles of PCR for library amplification. Sequencing data were generated on a NovaSeq 6000 using the paired end 150 cycle protocol.

### ATAC-seq data generation

For ATAC sequencing, we followed the protocol established by Buenrostro et al.^50^. Briefly, approximately 50,000 cells were used for nuclei preparation. Chromatin tagmentation and library construction were carried out using the Illumina Tagment DNA Enzyme and Buffer kit (Illumina). ATAC-seq DNA libraries were sequenced on the Illumina NextSeq 500 system with 76 bp paired-end reads.

### H3K27ac, H3K27me3, and MYB ChIP-seq data generation

ChIP-seq assays for H3K27ac (Abcam Cat #ab4729), H3K27me3 (Abcam Cat #ab192985), and MYB (Abcam Cat #ab45150) were performed using the following protocol. 40 million cells were used as the initial input per mark. The cells were pelleted by centrifugation at 1200g for 5 minutes (min) at room temperature (RT). The cells were resuspended in 1% formaldehyde (Thermo Fisher, 28906) in DPBS for 10 min at RT. The fixation was quenched with 0.225 M Glycine. The cells were then washed twice in DPBS with complete protease inhibitor (PI, Roche, 5892791001) and 1 mM final concentration of PMSF. After washing, the cells were resuspended in 5 ml covaris lysis buffer with PI. Cells were then washed and resuspended in covaris wash buffer with PI, and washed pellets were resuspended in covaris shearing buffer. The sample was transferred to a covaris millitubes (Covaris, 520135). The samples were sonicated with the following settings bath temp 6 °C, Duty Cycle 30%, Peak Incident Power 420, Cycles/Burst 200, for 12 min. The sonicated samples were used or stored at −80 °C until used. ChIP enrichment was performed using 60 μL of DiaMag A/G beads per 10 million cells. The samples were then precleared at 4°C for 1 hour. The antibody for the respective mark was then added to the sample and incubated with rotation at 4 ^°^C for 16-18 hours. The ChIP complex on beads was washed with DiaMag bead wash buffers: Low Salt Wash Buffer, High Salt Wash Buffer, LiCl Wash Buffer, and TE Buffer. The ChIP DNA was eluted using elution buffer (0.1 M NaHCO3, 1% SDS). The samples were reverse crosslinked at 65 ^°^C with shaking overnight. The ChIPed DNA was cleaned using the Qiagen MinElute kit according to the vendor’s instructions. The resulting DNA was quantified using the Agilent 4400 tapestation. Libraries were generated using the Kapa Hyper Prep Kit (Roche) according to the manufacturer’s instructions. Sequencing data were generated on a NovaSeq 6000 using the single-read, 100-cycle protocol.

### GATA3 and RUNX1 ChIP-seq data generation

10 million cells were crosslinked with 1% formaldehyde for 10 minutes at room temperature with gentle shaking. The reaction was quenched with 0.125 M glycine (Boston Bioproducts, Cat #C-4375) for 5 minutes at room temperature, and then the cell pellets were washed twice with PBS. The cells were stored at −80 °C until used. Sonication was performed using the Covaris M220 sonicator to generate chromatin fragments ranging from 200 to 700 bp. Chromatin immunoprecipitation was conducted using the Anti-RUNX1/AML1 antibody (Abcam Cat #ab272456) and GATA-3 (Cell Signaling Antibody Cat #5852). Diagenode iDeal ChIP-seq Kit for Transcription Factors (Cat # C01010054) was used according to the manufacturer’s protocol for the pull-down. Input controls (sonicated extracts) were included to support proper downstream bioinformatics analysis. Sequencing libraries were prepared with the KAPA HyperPrep Kit (Roche Cat # 07962347001) and indexed using Illumina TruSeq adapters (REF 20042113). The libraries were then cleaned, double-stranded selected, and sequenced on an Illumina NextSeq 2000.

### Hi-C data generation for PDX samples, and Jurkat and DND41 cell lines

Hi-C data was generated using the protocol as described by Rao et al^51^. Briefly, the samples were cultured under recommended conditions to about 80% confluence. Five million cells were crosslinked with 1% formaldehyde for 10 min at room temperature, then digested with 125 units of MboI, and labeled by biotinylated nucleotides and proximity ligated. After reverse crosslinking, ligated DNA was purified and sheared to 300–500 bp, then ligation junctions were pulled down with streptavidin beads and prepared as general Illumina library. The Hi-C samples were sequenced paired-end 151 cycles on Illumina Hiseq 4000.

### PacBio Iso-seq data generation for PDX samples and Jurkat cell line

Total RNA from the Jurkat cells and mouse PDX samples were extracted using the TRIzol^TM^ Reagent (ThermoFisher Scientific). RNA was subjected to DNAse I treatment and then purified with RNA Clean & Concentrator^TM^ −5 (Zymo Research). Iso-Seq libraries were prepared following the “Procedure & checklist - Preparing Iso-Seq libraries via SMRTbell prep kit 3.0” (PacBio). The final Iso-seq libraries were sequenced either on the PacBio Sequel system (for Jurkat cells) or the Sequel IIe system (for PDX samples).

## Analytical methods

### Samples from the T-ALL cohort and their annotation

For the analysis including T-ALL cohort, we used samples from the recent study^12^ and their annotations from the supplementary tables. The coordinates were converted to hg19 reference using LiftOver tool from ucsc v051223^52^ package. Subtype annotation was obtained from the Table S3^12^ (column “Reviewed.subtype”), and *TAL1* alterations were taken from the Table S9^12^, requiring column “Targeted.gene” to be *TAL1*. To identify alterations associated with the different known mechanisms of *TAL1* activation, we used column “Feature.(curated.name)”. For *TAL1* alterations annotation, the following features were selected (the annotations in the brackets correspond to the Feature.(curated.name) from Table S9 from^12^): TAL1 Downstream Enh I (’TAL1 Enhancer Gain Downstream I’ and ’TAL1 Enhancer SNV/Indel Downstream I’); TAL1 Downstream Enh II (‘TAL1 Enhancer SNV/Indel Downstream II’), TAL1 Intron Enh (‘TAL1 Enhancer SNV/Indel Intron’), TAL1 Upstream Enh (‘TAL1 Enhancer SNV/Indel Upstream’), STIL-TAL1 (STIL-TAL1). All other *TAL1* associated features were intra- and inter-chromosomal translocations and were included in the group ‘*TAL1* Translocation’, except those for 3 samples (SJALL015626_D, SJALL069889_D and SJALL071262_D for the following reasons: SV in the sample SJALL069889 involved exons of TAL1 (INV chr1_47225194 chr1_47985048), short SV in SJALL015626_D involved intron of TAL1 (chr1_47222950 chr1_47223055) and a short chr1 inversion in SJALL071262_D (INV chr1_45588233 chr1_47251827) annotated as “TAL1 Intergenic Inversion”. Sample SJTALL002048 didn’t have alterations associated with TAL1 listed in Table S9, though it had its reviewed subtype driver was annotated as “TAL1 enhancer SNV/Indel”, and based on the WGS analysis, the respective alteration was in the *TAL1* downstream enhancer I, so this sample was added to the *TAL1* Downstream Enh I group. Samples with *TAL1* features reported in the Table S9^12^ but not annotated as either TAL1 DP-like or TAL1 alpha/beta-like subtypes, were removed from the analysis, resulting in 404 samples.

Furthermore, samples that had more than one *TAL1* associated features (5 samples) were filtered out. Then, we carefully manually inspected the alterations associated with *TAL1* enhancers and found few samples with alterations located far away from the mutational hotspots in those groups (see **Extended Data Fig. 1** and **Supplementary Table 1**). Based on this procedure, 5 samples were filtered out: one from the *TAL1* downstream enhancer I; one sample from downstream enhancer II, one sample from intron enhancer and two samples from upstream enhancer group. Finally, for all *TAL1* downstream enhancer I samples we performed manual review of its alterations, using somatic WGS BAM files. During manual review, we observed that all variants should be classified as either ITDs or indels (see **Supplementary Table 1**). Additionally, 3 samples with ITDs were moved to *TAL1* downstream enhancer II group and one sample (SJALL069999_D) with indel (Indel chr1_47203728) was moved to *TAL1* downstream enhancer I group based on the coordinate of the alteration. As result, we had retained 391 samples in total: 17 samples with ITDs and 8 samples with indels in the *TAL1* downstream enhancer I group; 3 samples with ITDs, 6 samples with indels, and 6 samples with SNVs in *TAL1* downstream enhancer II group; 28 samples with SNVs in *TAL1* intron enhancer group; and 58 samples with indels in *TAL1* upstream enhancer group (**Extended Data Fig. 1** and **Supplementary Table 1**).

### WGS data processing and variant calling

SJTALL002048 and SJALL015708 paired tumor and normal WGS data were obtained from the previous study^11^.The data were mapped to reference human genome assembly GRCh37-lite with bwa^53^. Somatic variants in each tumor were analyzed by Bambino^54^ (SNV/Indel), CONSERTING^55^ (in both paired and tumor-only mode for somatic and germline copy number alterations) and CREST^55^ (for structural variations). The detected somatic aberrations were manually curated to further rule out false discoveries.

### scRNA-seq analysis from normal human bone marrow and thymus

We used scRNA-seq data from normal human bone marrow and thymus from Cordes et al. 2022^26^. Annotated RDS object for the integrated normal bone marrow and thymus data was obtained from GSE195812. Cell type annotation was used from meta data column ’Global Annotations’. DotPlot function from Seurat v5.1.0^56^ package was used to visualize relative RNA-seq expression among the cell types for *TAL1*, *CD34* and other gene markers.

### Hematopoiesis ATAC-seq data analysis

We used human bone marrow and peripheral blood ATAC-seq data of normal sorted cell populations: HSCs, MPPs, LMPPs, CLPs, CD4 and CD8 T-cells from Corces et al. 2016^28^. Data were visualized in UCSC genome browser using Track Hub, https://s3-us-west-1.amazonaws.com/chang-public-data/2016_NatGen_ATAC-AML/hub.txt. Capture Hi-C data for CD34+ cells were obtained from the previous study^34^ via https://3dgenome.fsm.northwestern.edu (accessed in June 2025).

### TFBS search

Position frequency matrices (PFMs) were obtained from the JASPAR2024 vertebrates core redundant database^57^ (https://jaspar.elixir.no/download/data/2024/CORE/JASPAR2024_CORE_vertebrates_redundant_pfms_meme.txt) and H13CORE PFMs from HOCOMOCO^58^ (https://hocomoco13.autosome.org/downloads_v13). Motif scanning was performed using FIMO from the MEME v5.5.4^59^ package with parameter --thresh 0.05 for p-value, and all others as default. To scan motifs in the sequences with indels from 8 downstream enhancer I samples, we used the indel sequence +/- 10bp flanking sites. We additionally filtered the resulting hits using the following motifs in the matched sequences based on the literature and PWMs from JASPAR2024 and HOCOMOCO databases: AGATA or TGATA for GATA3 (based on GATA3.H13CORE.1.SM.B and MA0037.1 from HOCOMOCO and JASPAR2024, respectively, and Tanaka et al.)^60^, NNCNGTTN for MYB (based on Ciciro et al.^61^ and MYB.H13CORE.0.P.B and MA0100.1 from HOCOMOCO and JASPAR2024, respectively), TGTGGTT for RUNX1 (based on Bowers et al.^62^ and RUNX1.H13CORE.0.P.B and MA0002.1 from HOCOMOCO and JASPAR2024, respectively), and CAGNTG for TAL1 (based on Gould et al.^63^ and TAL1.H13CORE.2.P.B and MA0091.2 from HOCOMOCO and JASPAR2024, respectively). The majority of the TFBSs were identified by using both HOCOMOCO and JASPAR2024 databases (**Supplementary Table 2**).

### PDX ChIP-seq data processing

For H3K27ac, H3K27me3, H3K4me1 and MYB read data were single-end, and for GATA3 and RUNX1 read data were paired-end, so the corresponding processing was done in the single-end and paired-end modes, respectively. ChIP-seq reads were first trimmed with trim-galore v0.6.10 and then aligned to the human genome (GRCh37-lite) with bwa mem^53^ (v0.7.12), then ambiguously-mapped and duplicate reads were removed. Fragment length was estimated based on a cross-correlation analysis by SPP v1.10.1^64^. Duplicates were marked with bamsormadup from biobambam v2.0.87 package^65^ and only non-duplicated reads have been kept by samtools v1.9^66^ (with parameter “-q 1 -F 1024” for single-end read data and “-q1 -F 1036” for paired read data to remove singletons). For merging BAM-files across replicates samtools v1.9^66^ merge was used. bedGraph files were generated via bedtools v2.31.0 genomecov function using scalefactor of 100,000,000/total mapped reads. Finally, we used bedGraphToBigWig function from ucsc v051223^52^ to convert bedGraph to BigWig track.

For peak calling, MACS2 v2.1.1^67^ was used. We further removed peaks from the Y chromosome, as well as those overlapping genomic regions containing ‘N’. All peaks were resized to 501 bp centered at the peak summit defined by MACS2. Enrichment heatmaps of co-accessible sites were made using deeptools v3.5.4 package^68^.

### Public ChIP-seq datasets processing

Public ChIP-seq data were obtained from the previous studies^15,22,69^ (see **Supplementary Table 5** for GEO ids). For Jurkat MYB, H3K27ac, CTCF, GATA3 and RUNX1, we used bowtie2, which allows insertions/deletions relative to the target genome, with parameters --rfg 1,1 -k 1^70^. For all other public ChIP-seq samples bwa aln^53^ was used. Fragment length was estimated based on a cross-correlation analysis by SPP v1.10.1^64^. Duplicates were marked with bamsormadup from biobambam v2.0.87 package^65^ and only non-duplicated reads have been kept by samtools v1.9^66^ (with parameter “-q 1 -F 1024”). bedGraph files were generated via bedtools v2.31.0 genomecov function using scalefactor of 100,000,000/total mapped reads. Finally, we used bedGraphToBigWig function from ucsc v051223^52^ to convert bedGraph to BigWig track.

### PDX ATAC-seq data processing

The reads were trimmed with trim-galore v0.6.10 with a paired-end mode and aligned to human genome hg19 (GRCh37-lite) by bwa mem v0.7.17^53^, duplicated reads were then marked with bamsormadup from biobambam v2.0.87 package^65^ and only non-duplicated proper paired reads have been kept by samtools v1.9^66^ (with parameter “-q 1 -F 1804”), and center 80 bp of fragments then were extracted by bedtools v2.31.0^71^. For merging BAM-files across replicates samtools v1.9^66^ merge was used. bedGraph files were generated via bedtools genomecov function using scalefactor of 1,000,000,000/total mapped reads. Finally, we used bedGraphToBigWig function from ucsc v051223^52^ to convert bedGraph to BigWig track.

### Mono-allelic expression analysis

To evaluate whether *TAL1* is mono-allelically expressed, we first used heterozygous WGS SNPs identified in the patient samples SJTALL002048 and SJALL015708 (see Methods above). Next, to evaluate VAF of those variants in WGS and RNA-seq data, we ran bam-readcount v1.0.1 (https://github.com/genome/bam-readcount) with parameters -q 10 -b 15 on the WGS and RNA-seq BAMs from tumor samples. We filtered out variants that had a total coverage in RNA-seq < 5 reads in total.

To evaluate coverage of SNPs in ChIP-seq data, we again used bam-readcount v1.0.1 with parameters -q 10 -b 15 using input germline variants identified in the WGS data, that were observed in the region downstream of *TAL1*. We filtered out variants that had a total coverage in ChIP-seq < 5 reads in total (**Supplementary Table 3**).

For haplotype phasing, GATK HaplotypeCaller v4.1.8.0 was applied to the normal BAM files of SJTALL002048 and SJALL015708 at the default settings. After lifting over the output VCF file to GRCh38, we generated a phased VCF file using SHAPEIT4 v4.2.1^72^ along with the 1KG reference panel (GRCh38) publicly available at https://ftp.1000genomes.ebi.ac.uk/vol1/ftp/data_collections/1000G_2504_high_coverage/working/20201028_3202_phased/. The coordinates were converted to hg19 reference using LiftOver from ucsc v051223^52^ package.

To calculate allelic imbalance, we performed a two-sided binomial test with a null probability of success 0.5 in a Bernoulli experiment. The resulting P-values were adjusted using Benjamini–Hochberg FDR method. Allelic imbalance events were called significant if they reached FDR < 0.05 and denoted as suggestive if they had 0.05 ≤ FDR < 0.15.

### Allele-specificity of TAL1 CRC binding sites

To calculate the number of ChIP-seq and ATAC-seq reads containing indels for SJTALL002048_X and Jurkat cell line samples, we used indelPost tool^73^, that performs read re-alignment for complex indels and outputs the counts of wild type and indel-containing reads with improved accuracy. For PDX sample, we used indel: chr1:47669403 TTATCAAAAAGACAGAAAA>T, and for Jurkat cell line we used indel: chr1:47704967 A>ACCGTTTCCTAAC as inputs for indelPost. Significance was evaluated using two-sided binomial test with a null probability of success 0.5 in a Bernoulli experiment, and the resulting P-values were adjusted with the FDR method.

### PacBio Iso-seq analysis

PacBio iso-seq data were analyzed using smrttools v13.0 (https://isoseq.how). First, primer removal and demultiplexing were performed with lima with the following parameters --isoseq --dump-clips --peek-guess. Then isoseq refine with parameter --require-polya was used for trimming polyA tails and concatemer removal. After that isoforms were called using isoseq cluster with parameter --use-qvs and then alignment to the hg19 (GRCh37-lite) was performed using pbmm2 align with parameters --preset ISOSEQ --sort --log-level INFO.

Then to collapse redundant transcripts to unique isoforms, isoseq collapse was run with parameter --do-not-collapse-extra-5exons. To classify and filter full-length transcript isoforms, pigeon workflow was used. First, pigeon prepare was used to sort transcript .gff (output from the “isoseq collapse” step). Then, pigeon classify with parameter --fl was used perform isoform classification. Finally, pigeon filter with parameter --isoforms was used to filter isoform, and then pigeon report was used to obtain gene saturation. bedGraph files were generated via bedtools v2.31.0 genomecov function using scalefactor of 1,000,000/total mapped reads. Finally, we used bedGraphToBigWig function from ucsc v051223^52^ to convert bedGraph to BigWig track.

### Hi-C analysis

The reads were trimmed with trim-galore v0.6.10 with a paired-end mode. Then trimmed reads were mapped to hg19 (GRCh37-lite) and processed using Juicer v1.6 ^74^ with default parameters and using bwa v0.7.17^53^. For PDX models, Hi-C experiment was performed on two replicates per each PDX. For those samples merged_nodups.txt files were merged using mega.sh from the juicer pipeline, and then .hic files were generated. The resulting .hic files with minimum mapping quality of 30 were visualized via GenomePaint^75^.

### RNA-seq analysis

RNA-seq data were mapped with STAR v2.3.0e_r291^76^. Gene-level read counts were generated with HTseq-count v0.11.2^77^, and the number of transcripts per million mapped reads (TPMs) were calculated based on the transcript models in GENCODE v19.

For merging BAM-files across replicates samtools v1.9^66^ merge was used. bedGraph files were generated via bedtools v2.31.0 genomecov function using scalefactor of 100,000,000/total mapped reads. Finally, we used bedGraphToBigWig function from ucsc v051223^52^ to convert bedGraph to BigWig track.

Processed RNA-seq data for 1,335 T-ALL cases (including 26 cases that lack WGS data) were used from the previous study of Polonen et al.^12^ and TPM matrix was obtained from Synapse under accession number syn54032669.

### DEG analysis

For the subtype marker DEG analysis, we compared samples from TAL1 DP-like subtype vs samples from TAL1 alpha/beta-like subtype as annotated in the column “Reviewed.subtype” in the Table S3 from the Polonen et al.^12^. Significance was evaluated using two-sided Wilcoxon rank-sum test, and P-values were adjusted using Benjamini–Hochberg FDR method.

To identify markers of the *TAL1* downstream enhancer I group compared to all other samples with other TAL1 alteration, we used linear model with gene expression as response variable, while presence in the *TAL1* downstream enhancer group and subtype were provided as explanatory variable and covariate, respectively. Benjamini–Hochberg FDR method was used for P-value adjustment.

### *TAL1* enhancer variants effects prediction with AlphaGenome

To predict *TAL1* variant effects we run AlphaGenome^23^ tool through the application programming interface (API) at http://deepmind.google.com/science/alphagenome. For this analysis we used variants affecting one of the four *TAL1* enhancers from 122 patients, which had variant REF/ALT information in the Table S14 of Polonen et al^12^. Sample SJALL069731_D with the largest ITD of >10 kbp length was excluded from this analysis. In total in this analysis, we used: 24 samples with variant in downstream enhancer I (16 ITDs and 8 indels), 14 samples with variants in downstream enhancer II (3 ITDs and 11 SNVs or indels), 28 samples with SNV variants in intron enhancer, 56 samples with indel variants in upstream enhancer. Some samples from downstream enhancer II, intron enhancer and upstream enhancer groups, had the same variants and as a result received the same predicted TAL1 expression scores, and so they were shown as single data points in **Fig. 5a** and **Extended Data Fig. 10a**. In particular, all SNV variants in the *TAL1* intron enhancer were the same (chr1: 47,696,311: C->T, hg38-based coordinates), and so they all received the same *TAL1* expression score of 0.04 by AlphaGenome.

For this analysis we used ontology of CD34+ common myeloid progenitors (CMPs, ontology code CL:0001059), the same ontology that was used by the published AlphaGenome study^23^ for *TAL1* non-coding variants analysis. Predicted *TAL1* expression scores correspond to the predicted *TAL1* expression change (REF - ALT) in CD34+ CMPs and they were obtained from AlphaGenome output table (tal1_diff_in_cd34 column). The predictions were made using hg38 genome, and to obtain the coordinates of the corresponding region in hg19 for comparison with the experimental data, we used LiftOver tool from ucsc v051223.

### Enrichment of T-ALL driver gene mutations in *TAL1* samples groups

T-ALL driver gene mutations for samples from six *TAL1* groups were obtained from Table S9 of Polonen et al^12^. The oncogenes and tumor suppressor genes were selected if they had >10% mutation frequency in at least one of the six *TAL1* groups. To calculate driver mutation enrichment in each group vs samples in all other groups, we used Fisher’s exact test, and the resulted P-values were adjusted with Benjamini–Hochberg FDR method.

### Visualization of coverage tracks

For visualization of coverage tracks we used GenomePaint on the St. Jude Cloud (https://genomepaint.stjude.cloud/^75^).

## Extended 1116 Data Figure legends

**Extended Data Fig. 1:**
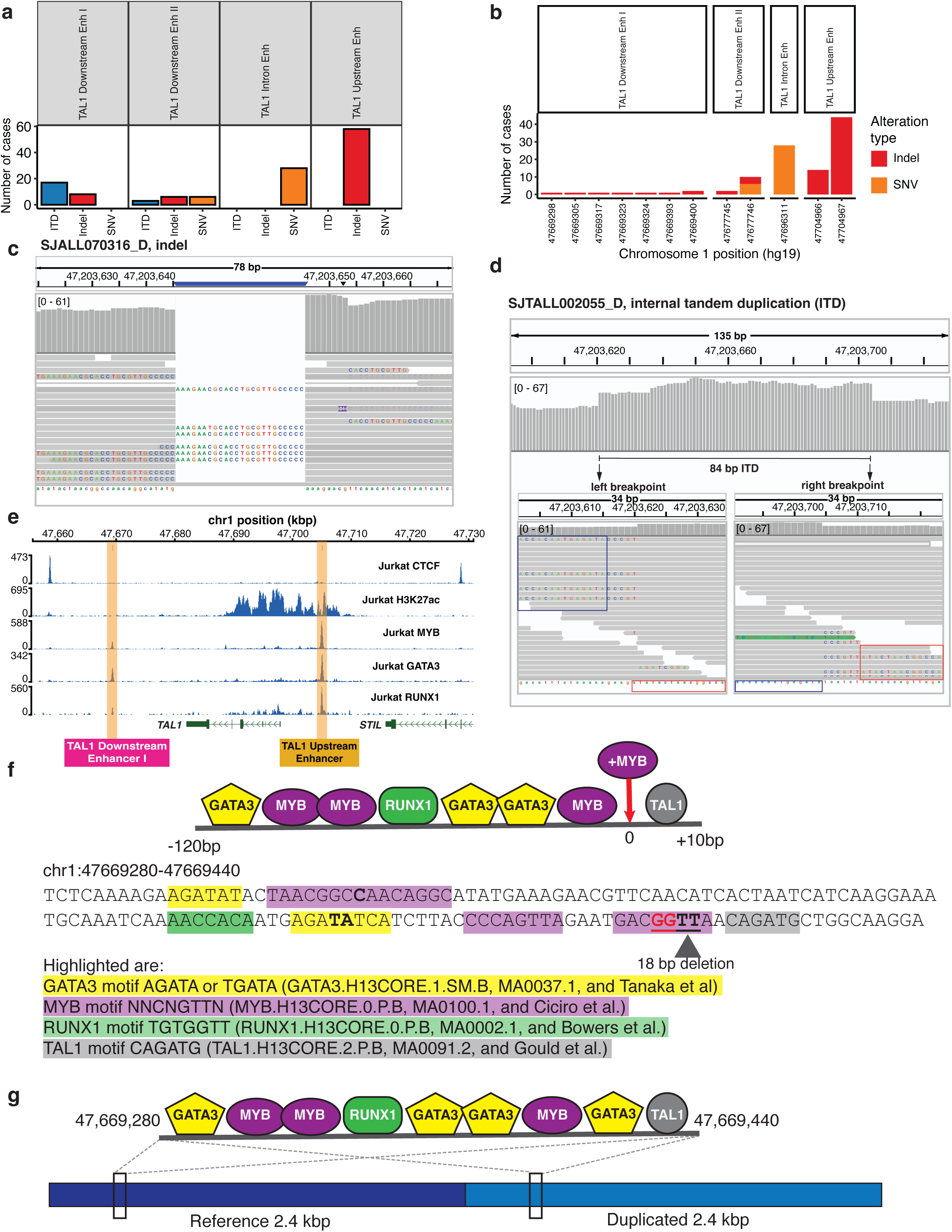
Somatic non-coding variants resulting in activated T-ALLs *TAL1* enhancers in four different regions. **a**, Barplot showing distribution of somatic ITD, indel and SNV alterations across the four enhancer regions in T-ALL samples. **b**, Recurrence of SNV and indel variants at distinct genomic loci in the four enhancer regions. **c-d**, Examples of manually curated and re-annotated *TAL1* alterations (note, the reads are mapped to hg38 reference genome). **c**, WGS reads of a heterozygous indel in the sample SJALL070316_D displayed using the IGV browser. Two alterations were originally reported in the region: INV chr1_47203645 chr1_47203646 and Indel chr1_47203645 (both in hg38 coordinates) in Polonen et al.^12^, and after manual review the alteration was annotated as indel (hg38 coordinates: Indel chr1_47203645 and lifted to hg19 coordinates: Indel chr1_47669317, and see **Supplementary Table 1d**). **d**, WGS reads of a heterozygous ITD in the sample SJTALL002055_D displayed using the IGV browser. Two alterations were originally reported in the region: Indel chr1_47203620 and INV chr1_47203621 chr1_47203704 (both in hg38 coordinates) in Polonen et al.^12^, and after manual review the alteration was annotated as ITD (hg38 coordinates: ITD chr1_47203621 chr1_47203704 and lifted to hg19 coordinates: ITD chr1_47669293 chr1_47669376, and see **Supplementary Table 1d**). **e**, Residual MYB, GATA3 and RUNX1 binding at downstream enhancer I region in Jurkat cell line based on ChIP-seq coverage plot. ChIP-seq data for CTCF, H3K27ac, GATA3 and RUNX1 were obtained from Hnisz et al. 2016^69^ and MYB ChIP-seq data was from Mansour et al. 2014^15^. The location of the *TAL1* upstream neo-enhancer caused by a somatic indel is highlighted along with the downstream enhancer I region where there is no somatic mutation in Jurkat cells. **f**, On top, schematic showing TAL1 CRC TFs, e.g. GATA3, RUNX1, MYB, and TAL1, that have motifs present in the reference genome as well as *de novo* MYB TFBS that was introduced by the complex indel in SJTALL002048 sample. At the bottom, sequence of a 160 bp region containing binding sites motifs of TAL1 CRC TFs. The complex indel site is shown underlined in bold. The overlapping bases for two MYB (#1 and #2) and for two GATA3 (#2 and #3) motifs are in bold. Of note, the 18 bp deletion removes reference GATA3 TF motif #4. **g**, Schematic showing TAL1 CRC TFs that have motifs present in the 160 bp region of the reference genome and its respective location in the duplicated 2.4 kbp region of the ITD sample SJALL015708.

**Extended Data Fig. 2:**
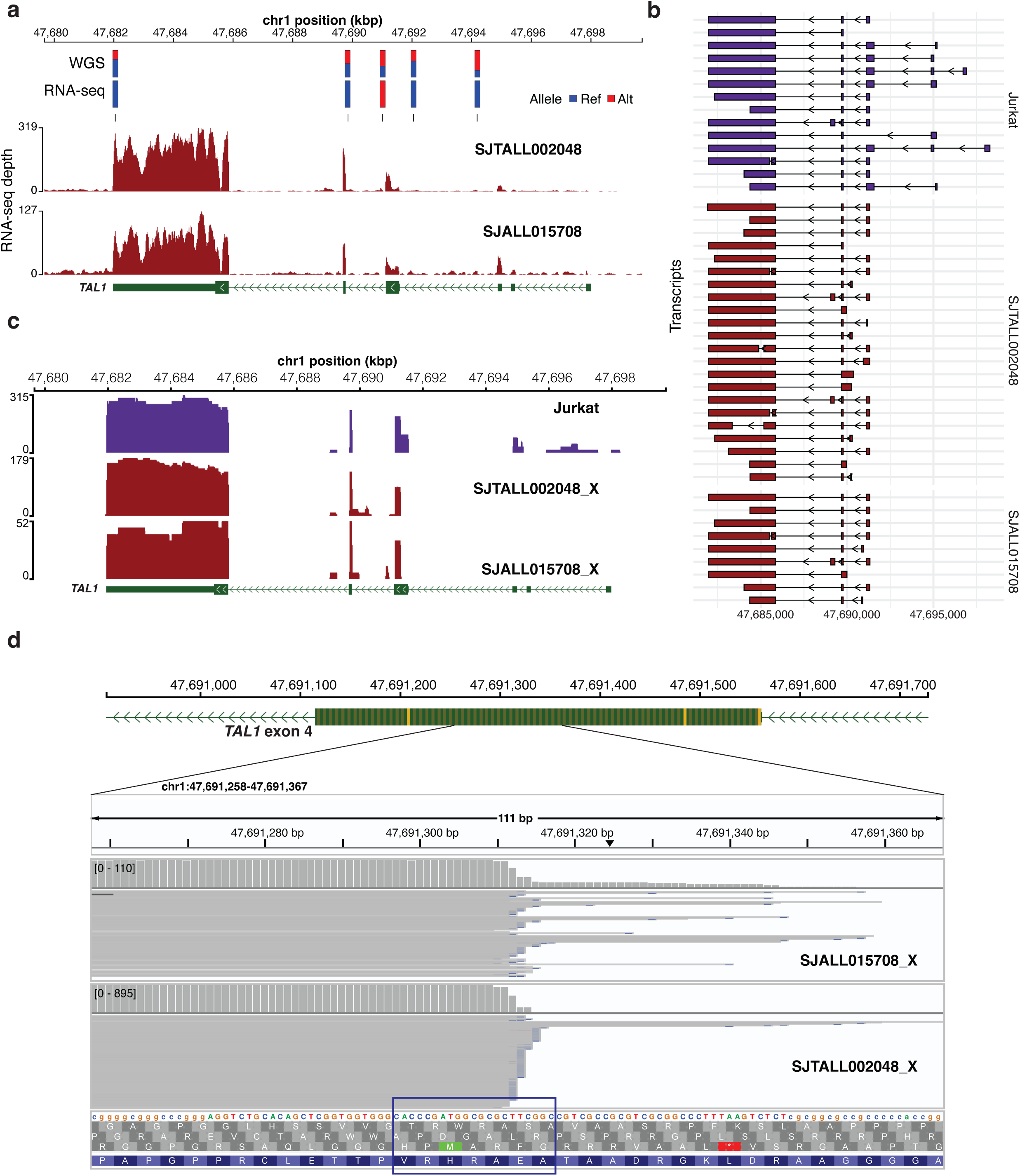
*TAL1* transcription in the two index samples SJTALL002048_D and SJALL015708 from downstream enhancer I group. **a**, Allele-specific expression of *TAL1* in RNA-seq data generated from the patient T-ALL samples. At the bottom, coverage plot showing RNA-seq expression. On top, barplots show variant allele frequencies (VAFs) of REF (reference) and ALT (alternative) alleles in patient tumor WGS and RNA-seq data of the indel sample SJTALL002048_D. Such analysis cannot be performed for the ITD sample SJALL015708 due to lack of heterozygous SNPs. **b**, Plot showing all transcripts obtained from PacBio Iso-seq for Jurkat cell line and two PDX samples. **c**, *TAL1* expression by PacBio Iso-seq full-length sequencing. The coverage plots of the two PDX samples (SJTALL002048_X and SJTALL015708_X) are shown along with that of Jurkat cell line. **d**, On top, the view of the TAL1 exon 4 from GenomePaint^75^, with translation initiation sites (ATG codons) highlighted in orange. At the bottom, transcription initiation in *TAL1* exon 4 from the PacBio Iso-seq reads in two PDX samples shown in IGV viewer. The blue rectangle shows the sequence corresponding to previously reported +1 promoter at exon 4 (from Fig. 1C in Courtes et al.^78^).

**Extended Data Fig. 3:**
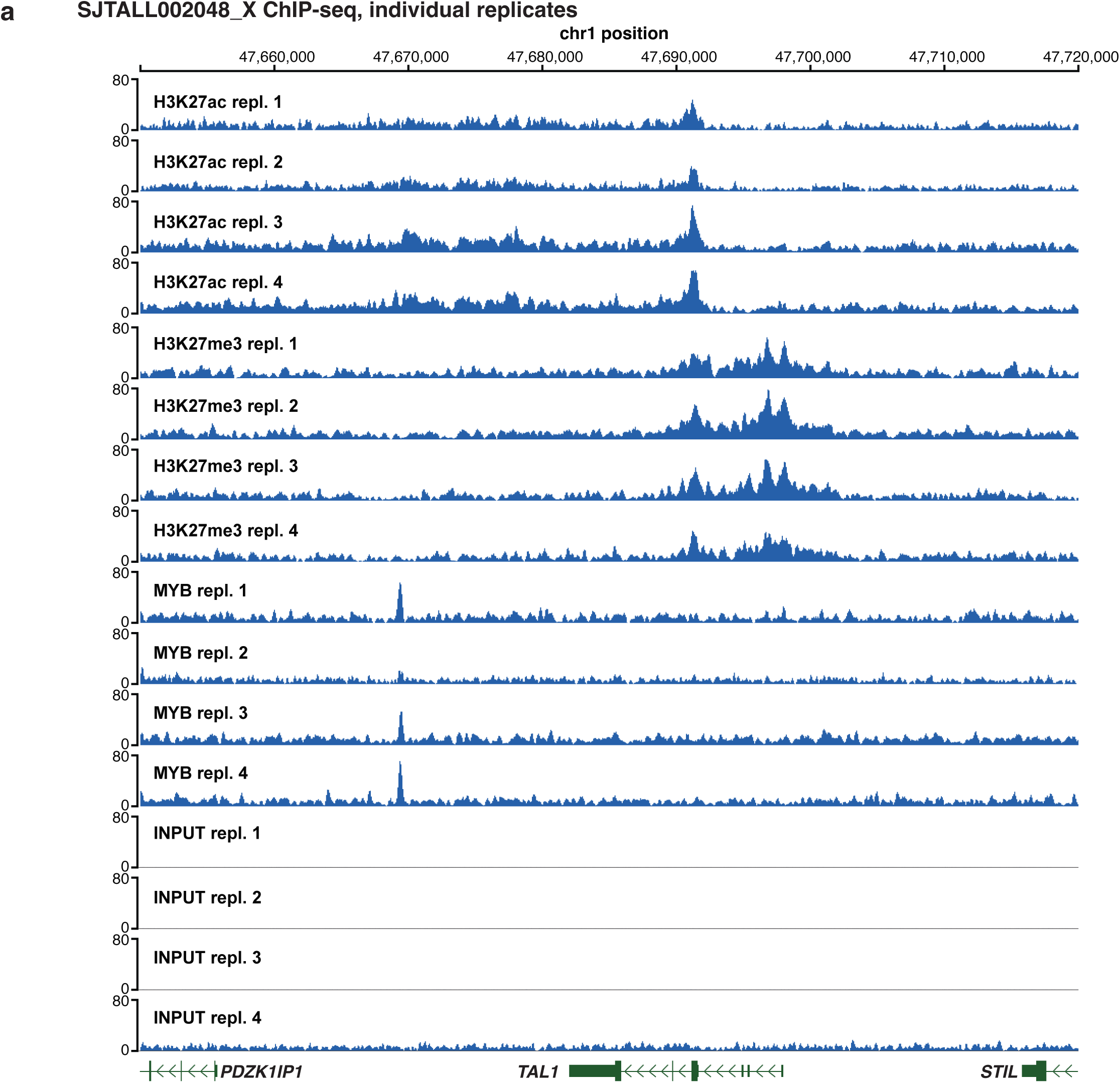
Replicates of ChIP-seq data in SJTALL002048_X PDX sample. Coverage plots of four individual ChIP-seq replicates for H3K27ac, H3K27me3, MYB antibodies and Input control. The scale is fixed to range from 0 to 80.

**Extended Data Fig. 4:**
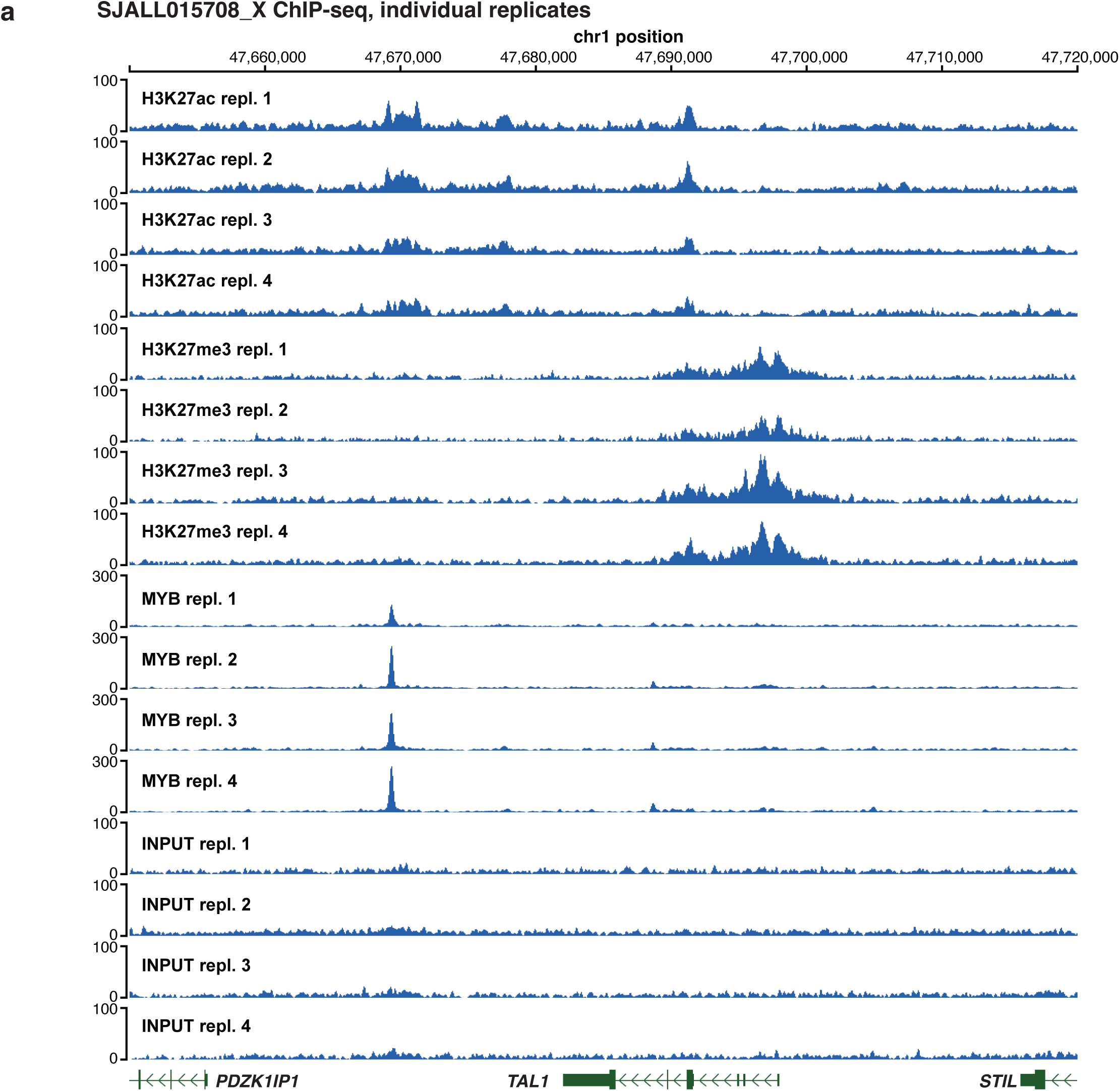
Replicates of ChIP-seq data in SJALL015708_X PDX sample. Coverage plots of four individual ChIP-seq replicates for H3K27ac, H3K27me3, MYB antibodies and Input control. The scale is fixed to ranges from 0 to 300 for MYB tracks and from 0 to 100 for all other tracks.

**Extended Data Fig. 5:**
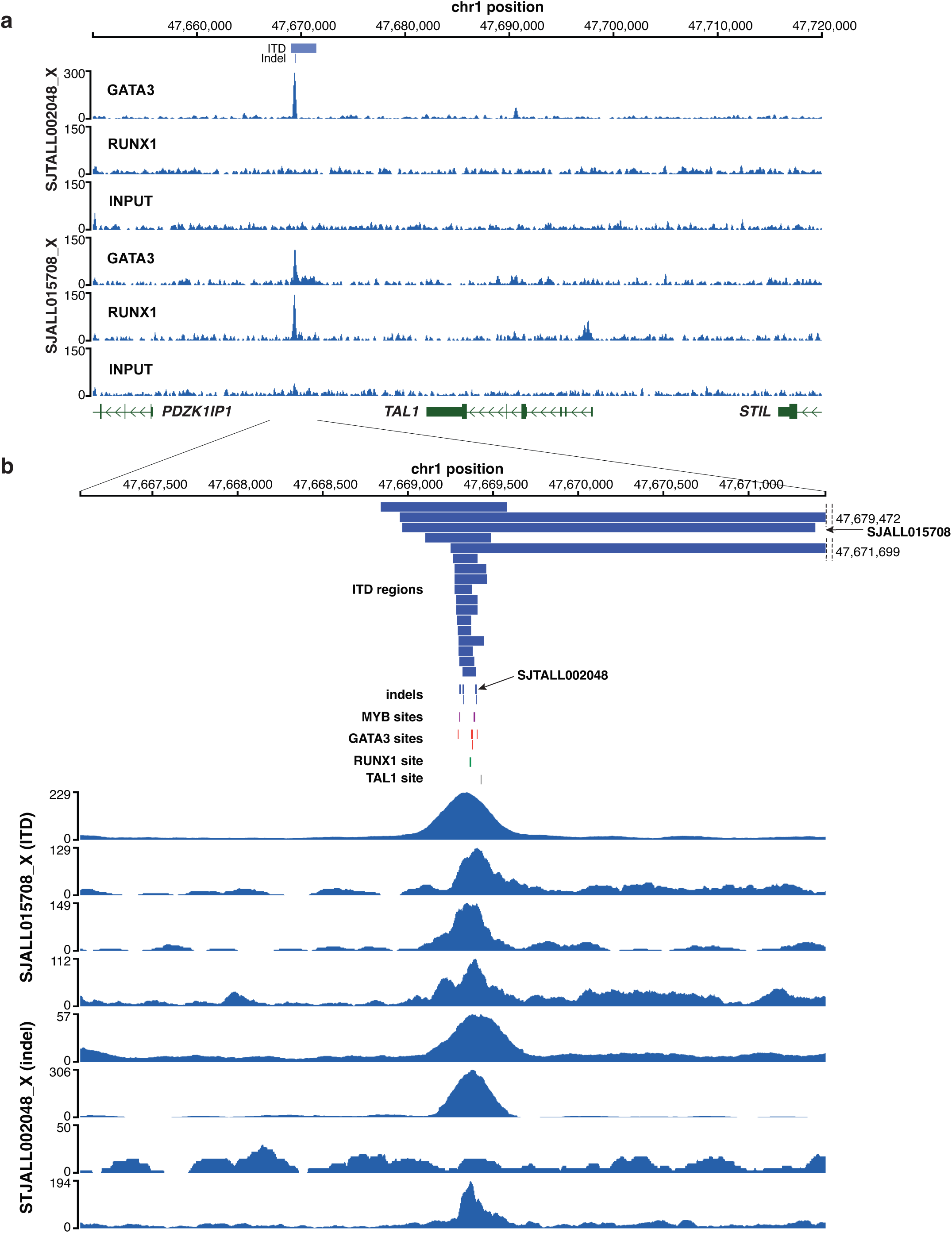
Recurrent TF ChIP-seq peaks in downstream enhancer I regions. **a**, Coverage plots of ChIP-seq for GATA3, RUNX1 and Input for SJTALL002048_X and SJTALL015708_X PDX samples. The scale is fixed to ranges from 0 to 300 for GATA3 track of SJTALL002048_X and from 0 to 150 for all other tracks. The region at the bottom corresponds to the location of the genomic segment shown in panel **b**. **b**, Alignment of ChIP-seq peaks in the two PDX samples to canonical TFBSs. On top, genomic locations of ITDs and indels in downstream enhancer I region with the labeled locations for the two index samples. In the middle, locations of canonical MYB, GATA3, RUNX1, and TAL1 TFBSs in the reference genome. At the bottom, coverage plots showing ChIP-seq peaks for MYB, GATA3 and RUNX1 antibodies, all of which are aligned to the canonical TFBSs despite the different locations of somatic alterations in these two samples.

**Extended Data Fig. 6:**
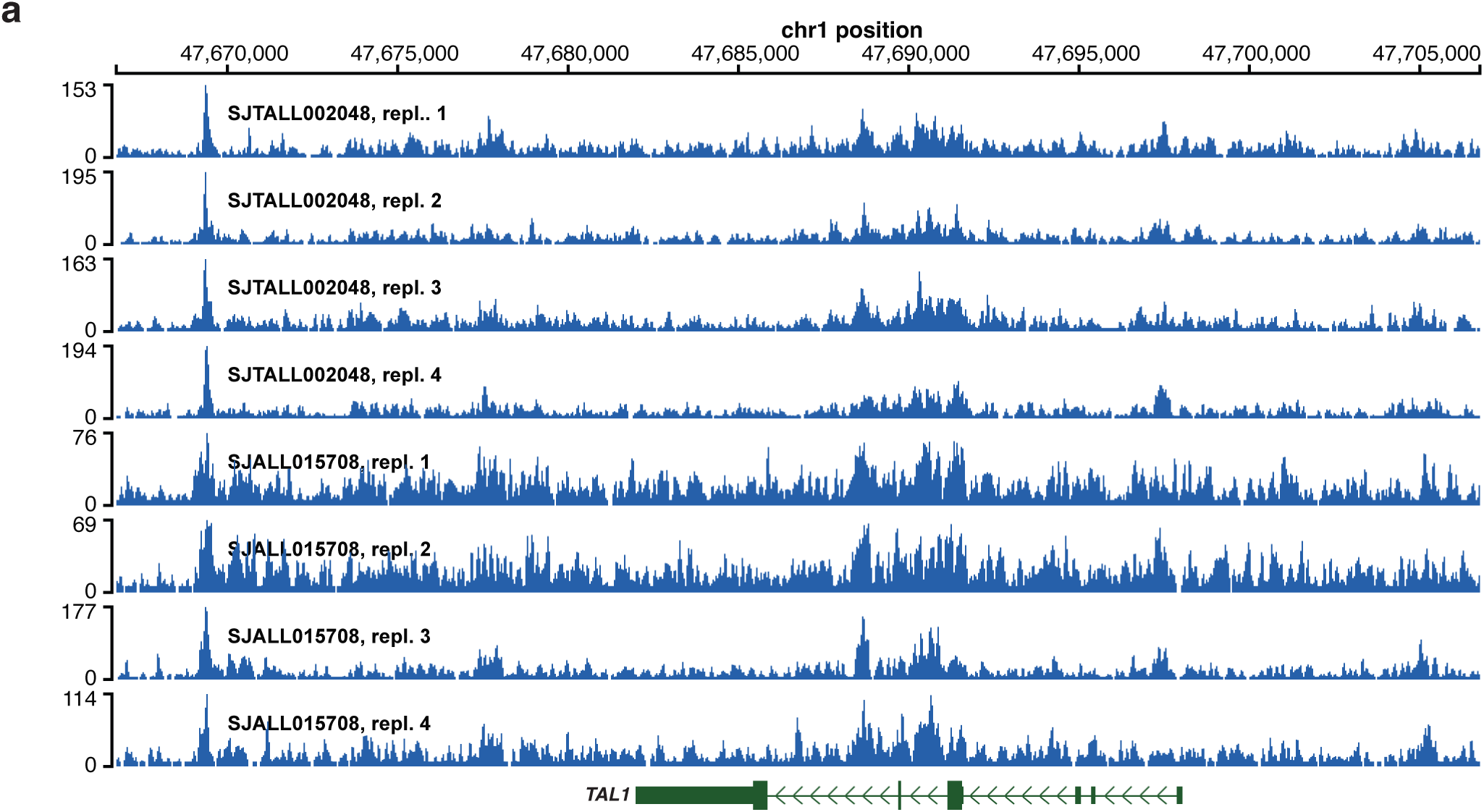
Replicates of ATAC-seq data in SJTALL002048_X and SJALL015708_X PDX samples. **a**, Coverage plots of four individual replicates of ATAC-seq for SJTALL002048_X and SJALL015708_X PDX samples.

**Extended Data Fig. 7.**
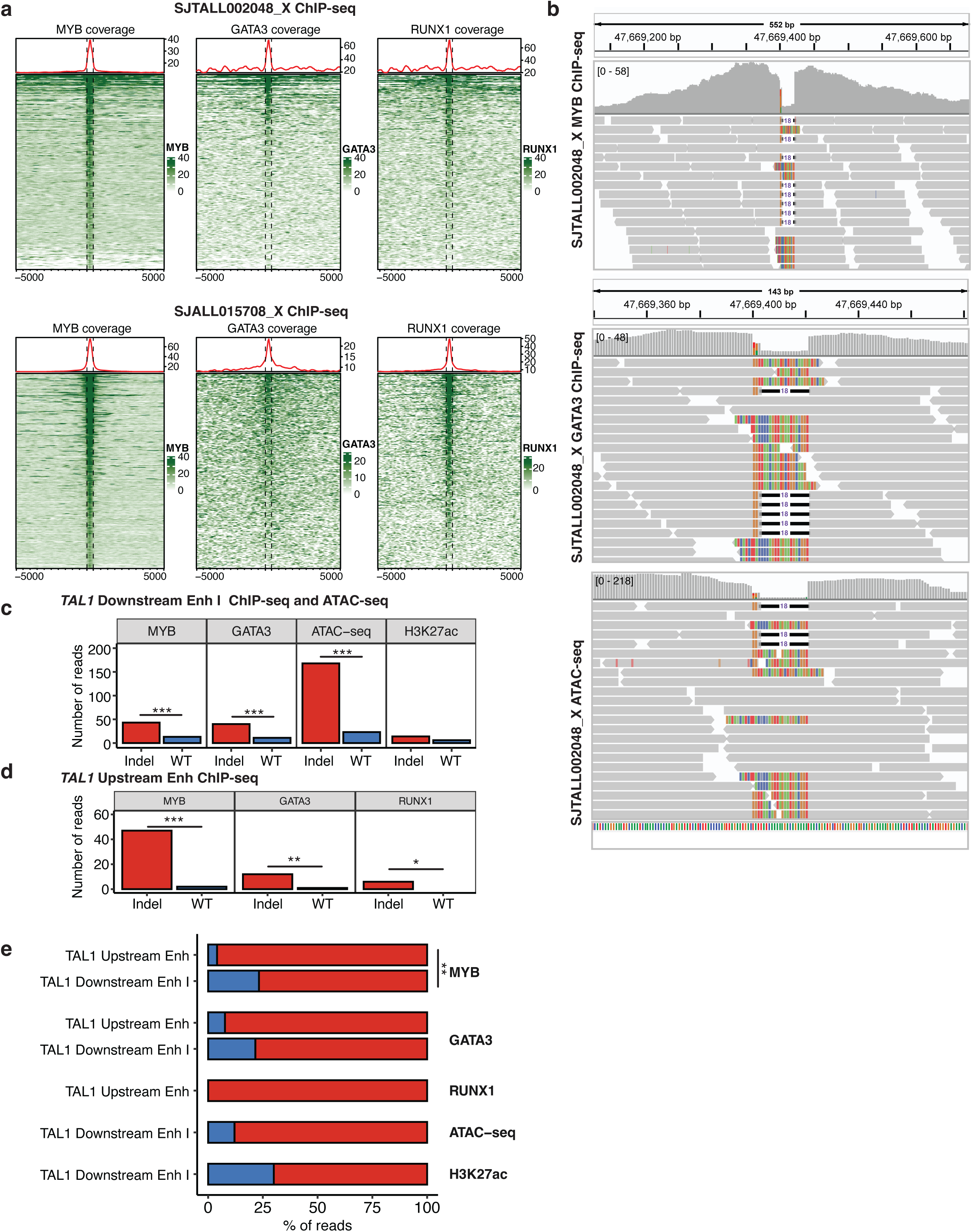
ChIP-seq profiles of PDX models derived from the two index samples. **a**, Heatmaps showing ChIP-seq coverage for MYB, GATA3 and RUNX1 antibodies centered around MYB peaks for the indel sample SJTALL002048_X (on top) and the ITD sample SJTALL015708_X (at the bottom). **b**, Wild type and mutant alleles of complex indel of SJTALL002048_X in MYB ChIP-seq (on top), GATA3 ChIP-seq (in the middle) and ATAC-seq (at the bottom) data displayed by IGV viewer. The majority of reads either contain indel or have soft-clipped sequence overlapping the indel region. **c-d**, Differential binding number of reads overlapping indel and WT alleles in ChIP-seq and ATAC-seq data of SJTALL002048_X sample (*TAL1* downstream enhancer I region, **c**) and data of Jurkat cell line samples (*TAL1* upstream enhancer, **d**). Indel and WT allele-containing read counts were obtained using the indel-post tool^73^. The significance was evaluated using two-sided binomial test, and the resulting P-values were adjusted with the FDR method; *** corresponds to FDR < 0.001, ** corresponds to FDR < 0.01, and * corresponds to FDR < 0.05. **e**, Percentage of reads overlapping indel and WT alleles in ChIP-seq of Jurkat cell line sample (*TAL1* upstream enhancer) and ChIP-seq and ATAC-seq of SJTALL002048_X sample (*TAL1* downstream enhancer I region). The significance was evaluated using Fisher’s exact test, and ** corresponds to P-value < 0.01.

**Extended Data Fig. 8:**
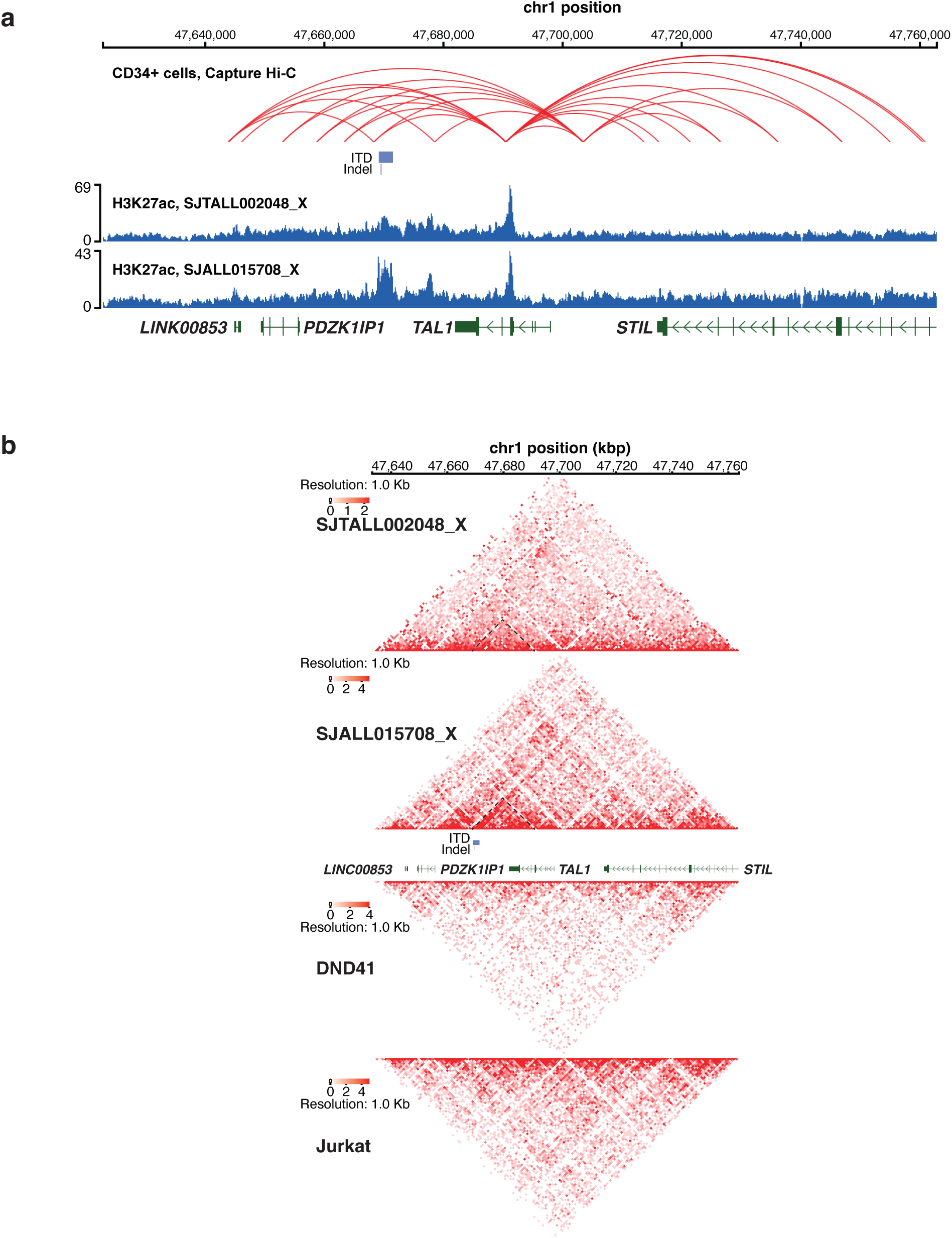
Chromatin contact between the downstream enhancers and *TAL1* promoter. **a**, Capture Hi-C in CD34 cells from the previous study^34^ which shows contact between downstream enhancer I region and exon 4 promoter. The contact regions also overlap with H3K27ac peaks from the two PDX samples shown at the bottom. **b**, Hi-C coverage of the region around *TAL1* locus in the two PDX samples (SJTALL002048_X and SJTALL015708_X), *TAL1*-negative cell line DND41 and *TAL1*-positive cell line Jurkat. The contact regions with enriched coverage in the two PDX samples are highlighted by dashed lines (forming triangles).

**Extended Data Fig. 9:**
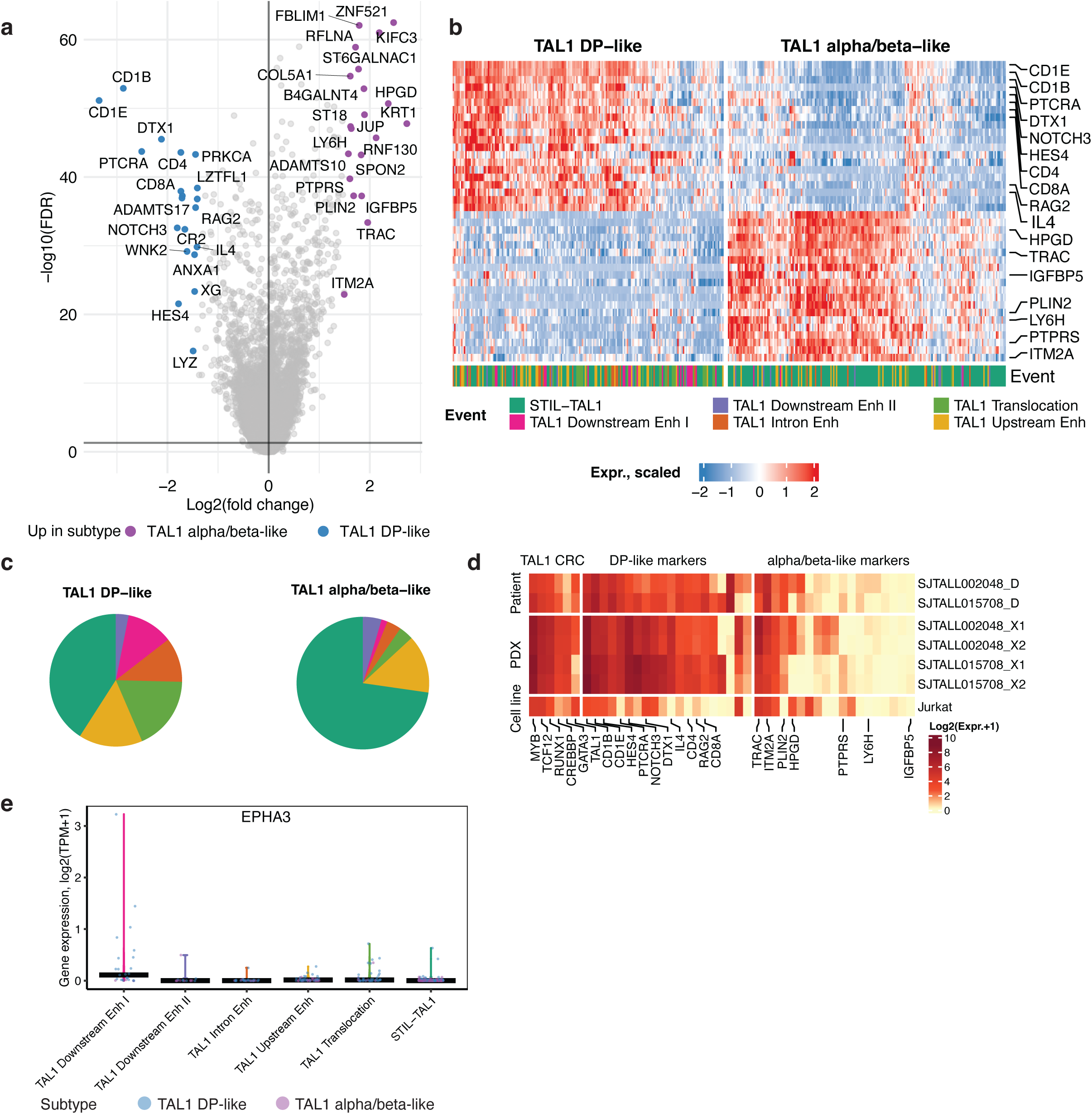
Expression profile of *TAL1* DP-like and alpha/beta-like subtypes. **a**, Volcano plot showing differentially expressed genes between *TAL1* DP-like and *TAL1* alpha/beta-like subtypes. Y-axis reflects the -log_10_ of the FDR adjusted Wilcoxon two-sided P-values. The top significant DEGs selected by fold change are highlighted and labeled for both subtypes. **b**, Heatmap shows RNA-seq expression of top markers between *TAL1* DP-like and *TAL1* alpha/beta-like subtypes from panel (A) in T-ALL samples. Each column represents one sample which is colored by its *TAL1* activation group at the bottom. **c**, Pie charts showing the composition of *TAL1* activation groups in DP-like and alpha/beta-like subtypes. The activation groups are color-coded using the same style as in panel **b**. **d**, Heatmap showing RNA-seq expression (log_2_(TPM+1) values) of TAL1 core regulatory circuit (CRC) genes (left) and top markers between *TAL1* DP-like and *TAL1* alpha/beta-like subtypes (right) for the following samples: SJTALL002048_D and SJTALL015708_D, two replicates of PDX samples: SJTALL002048_X and SJTALL015708_X, and Jurkat cell line. **e**, Violin plot showing RNA-seq expression (log_2_(TPM+1) values) of the *EPHA3* gene in the six groups with different *TAL1* activation mechanisms.

**Extended Data Fig. 10:**
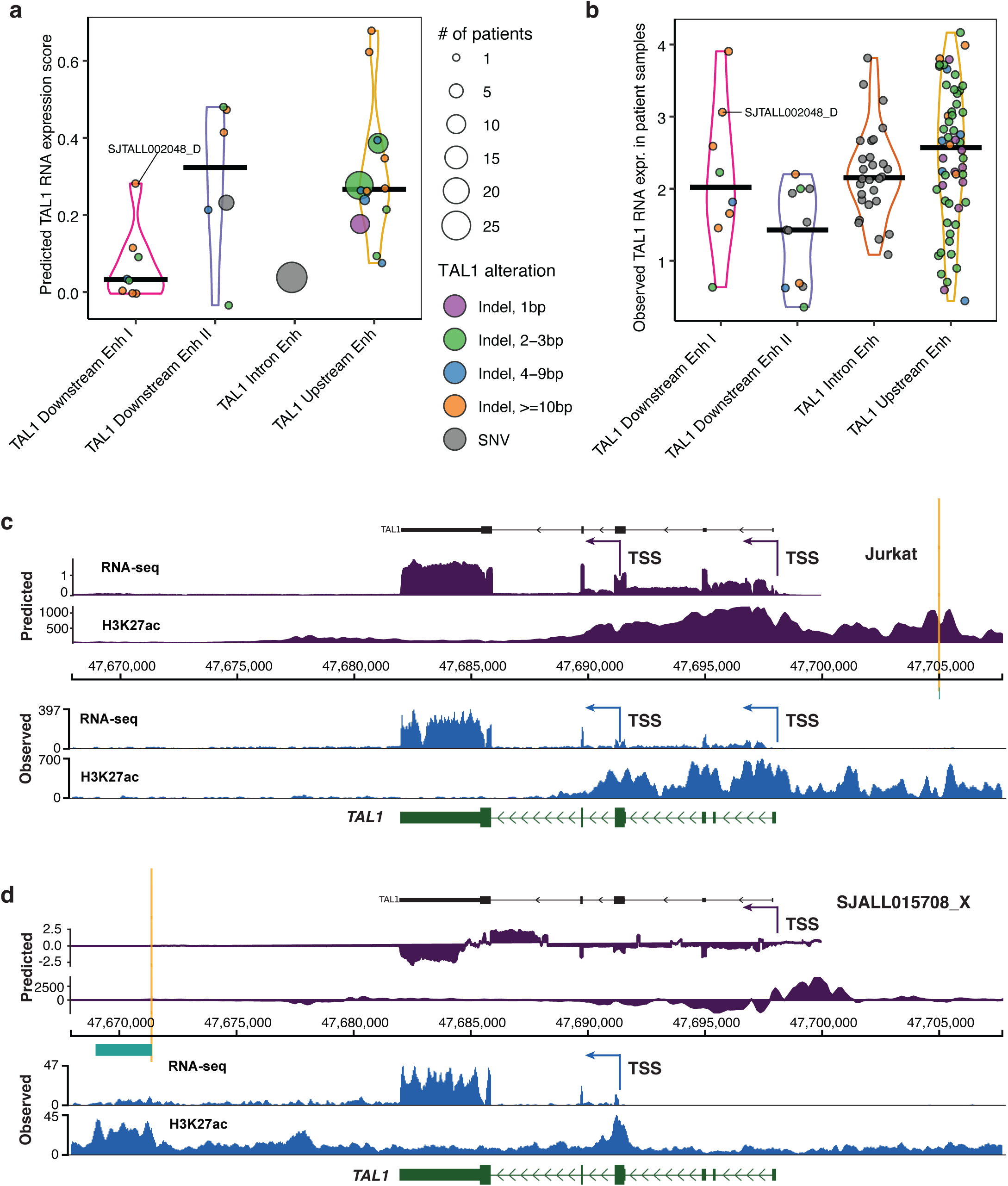
Evaluation of AlphaGenome predictions for effects of variants in *TAL1* enhancers. **a**, Violin pot showing distribution of the *TAL1* RNA expression scores predicted by AlphaGenome for the indel and SNV variants that affect one of the four enhancer regions mediating *TAL1* expression in 103 T-ALL patients. Dot size corresponds to variant recurrence, i.e. the number of T-ALLs with the same variant. **b**, Violin plot showing distribution of *TAL1* RNA-seq expression (log_2_(TPM+1) values) in 103 T-ALLs with variants affecting *TAL1* expression that are shown in panel **a**. **c-d**, Predicted and observed coverage tracks for *TAL1* RNA-seq and H3K27ac ChIP-seq in Jurkat cells with *TAL1* upstream enhancer mutation (**c**), and in SJALL015708_X case with ITD affecting *TAL1* downstream enhancer I (**d**). On top, coverage tracks show predicted by AlphaGenome *TAL1* RNA expression and H3K27ac ChIP-seq scores (difference between ALT and REF sequences). At the bottom, coverage plots show *TAL1* expression and H3K27ac ChIP-seq signal from the experimental data. Arrows mark the *TAL1* TSS at exon 1, and alternative *TAL1* TSS at exon 4.

## Supplementary Tables

**Supplementary Table 1: Samples with alterations that mediated *TAL1* activation. a**, Table containing annotation of 391 samples from six groups of *TAL1* alterations: *STIL*-*TAL1* deletion, *TAL1* translocations and four *TAL1* enhancers alterations (Final_annotation column). Reviewed.subtype contains subtype information from Polonen et al.^12^. previous_Feature(curated.name) column contains information about previous *TAL1* alteration annotation from Polonen et al. (corresponds to Feature(curated.name) from Table S9 of Polonen et al.). **b**, Table containing alterations in four *TAL1* enhancers that mediated its activation. The data were used for summary of alterations in four *TAL1* enhancers (**Extended Data Fig. 1a-b**). **c**, Table containing alterations in samples from *STIL*-*TAL1* and *TAL1* translocations groups. **d**, Table containing manually reviewed alterations in *TAL1* downstream enhancer I group of samples. “previous_variantID” columns contain information from Table S9 of Polonen et al., and “variantID” columns contain final variants after manual review (see Methods).

**Supplementary Table 2: TAL1, GATA3, RUNX1 and MYB TFBS mappings. a**, Table containing TAL1, GATA3, RUNX1 and MYB TFBS mappings (underlined) to the region of ITDs overlap. **b**, Table containing indel with +/-10 bp flanking sequences from 8 T-ALL samples with MYB TFBSs being shown underlined. **c**, Table containing MYB TFBS mappings (shown underlined) to the indel samples’ sequences from **Supplementary Table 2b**. **d**, Table containing ITD junction sequences, that include +/-10 bp flanking sequences and interstitial sequences (shown in red) if any. The “I” bar indicates the location of junction, upper case letters correspond to the left flanking sequence, while lower case letters correspond to the flanking sequence on the right. The underlined sequences correspond to the MYB TFBSs that were found. **e**, Table containing MYB TFBS mappings (shown underlined) to the indel samples’ sequences from **Supplementary Table 2d**. In **Supplementary Table 2a, c** and **e**, the results are shown for FIMO using HOCOMOCO and JASPAR2024 TF binding profiles databases.

**Supplementary Table 3: List of heterozygous SNPs with coverage in H3K27ac, H3K27me3 and PDX RNA-seq assays with haplotype assigned.** Table contains depth of reference (REF) and alternative (ALT) alleles in tumor and normal samples (obtained from bambino output^54^), as well as in the H3K27ac and H3K27me3 assays (see Methods). Column “haplotype” refers to whether the alternative allele is assigned to Hap1 (denoted 1) or Hap2 (denoted 2) based on haplotype phasing. “Assay_vaf” column corresponds to variant allele frequency in either of the H3K27ac, H3K27me3 or RNA-seq assays. In the downstream enhancer I and *TAL1* exon IV regions, Hap1 was found to be enriched in H3K27ac and RNA-seq, while Hap2 was found to be enriched in H3K27me3, and it can be seen that for sites with ALT allele assigned to Hap1, both H3K27ac and RNA-seq assays have higher “Assay_vaf” (>0.5), while H3K27me3 shows lower “Assay_vaf” (<0.5), and the pattern is reversed for the sites with ALT alleles assigned to Hap2. The read count of reference and alternative alleles in tumor and normal DNA samples are in columns N-Q and show balanced representation of these sites in DNA along with their population frequency in gnomAD database (“gnomadG2_AF” column). The significance of allelic imbalance of each site in the RNA-seq or ChIP-seq assays was calculated using two-sided binomial test with a null probability of success 0.5 in a Bernoulli experiment. The resulting P-values (“P-value” column) were adjusted using Benjamini–Hochberg FDR method (“FDR” column).

**Supplementary Table 4: DEG analyses results. a**, DEGs between DP-like and alpha-beta-like *TAL1* subtypes. DEGs were calculated using two-sided Wilcoxon test, and P-values were adjusted using the Benjamini–Hochberg FDR method. Table contains DEGs with FDR < 0.05. **b**, Expression profile of *TAL1* downstream enhancer I samples. Table containing DEGs between *TAL1* downstream enhancer I samples and samples with all other *TAL1* activating alterations used in the study. DEGs were calculated using linear model with *TAL1* subtype as a covariate, and P-values were adjusted using the Benjamini–Hochberg FDR method. Table contains DEGs with FDR < 0.25.

**Supplementary Table 5: List of ChIP-seq public datasets used in this study.** Table contains GEO ids and references of published ChIP-seq datasets used in the study.

## Data availability

Processed data for PDX ChIP-seq data for cell lines will be available from the Gene Expression Omnibus (GEO). Raw data for PDX models are being deposited to European Genome-phenome Archive (EGA). The two PDX models are available to the scientific community by writing to Dr. Keith L. Ligon.

## Code availability

indelPost tool that was used in the paper is available from the GitHub repository: https://github.com/stjude/indelPost. The scripts and additional information for reproducing analysis and figures are available at the GitHub page: https://github.com/nvterekhanova/TAL1_noncoding_publication.

## Acknowledgements

We thank the patients, staff, and scientists who contributed to this study, the Children’s Oncology Group (COG) operations staff, and the staff at the COG biorepository at Nationwide Children’s Hospital. We also thank Dr. Brian J. Abraham for critical review of the manuscript and Dr. Donald Yergeau who performed the RNA-seq data generation for Jurkat cell line. The study is supported by National Cancer Institute (NCI) grants R01CA216391 to J.Z., the St. Jude Research Collaboratives, the Consortium of the 3D Genome to J.Z. and A.T.L. All authors from St Jude received support from NCI Cancer Center grant P30 CA021765 and from American Lebanese Syrian Associated Charities (ALSAC). The AALL0434 clinical trial and AALL03B1 biobanking study were supported by NCTN Operations Center Grant (U10CA180886), NCTN Statistics and Data Center Grant (U10CA180889), COG Biospecimen Bank Grant (U24CA196173), and the St. Baldrick’s Foundation. The AALL0434 clinical trial was supported by Sandoz Inc.

## Author contributions

Study conception and design: J.Z. and A.T.L.; developed and performed experiments or data collection: J.E., S.N., A.T.L., D.T.T., K.-H. C., Y.S., L.D., B.J., V.V., H.Z., and X.M.; PDX models creation: A.T.L., K.-H. C., and D.T.T.; computation and statistical analysis: N.V.T., X.C., W.Y., Y.L., and K.H.; data interpretation, biological analysis and figure generation: N.V.T., J.Z., A.T.L., X.C., W.Y., Y.L., K.H., and X.M.; writing – original draft: N.V.T. and J.Z.; review and editing: J.Z., A.T.L., D.T.T., N.V.T., X.C., K.-H.C., B.J., V.V., S.N., and J.E.; administration: J.Z., A.T.L., and J.E. All authors read and approved the manuscript.

## References

1. Robb, L. et al. Absence of yolk sac hematopoiesis from mice with a targeted disruption of the scl gene. Proceedings of the National Academy of Sciences 92, 7075–7079 (1995).

2. Shivdasani, R.A., Mayer, E.L. & Orkin, S.H. Absence of blood formation in mice lacking the T-cell leukaemia oncoprotein tal-1/SCL. Nature 373, 432–434 (1995).

3. Porcher, C. et al. The T Cell Leukemia Oncoprotein SCL/tal-1 Is Essential for Development of All Hematopoietic Lineages. Cell 86, 47–57 (1996).

4. Robb, L. et al. The scl gene product is required for the generation of all hematopoietic lineages in the adult mouse. The EMBO Journal 15(1996).

5. Mouthon, M.-A. et al. Expression of tal-I and GATA-Binding Proteins During Human Hematopoiesis. Blood 81, 647–655 (1993).

6. Herblot, S., SteZ, A.-M., Hugo, P., Aplan, P.D. & Hoang, T. SCL and LMO1 alter thymocyte diZerentiation: inhibition of E2A-HEB function and pre-Tα chain expression. Nature Immunology 1, 138–144 (2000).

7. Delabesse, E. et al. Transcriptional Regulation of the SCL Locus: Identification of an Enhancer That Targets the Primitive Erythroid Lineage In Vivo. Molecular and Cellular Biology 25, 5215–5225 (2005).

8. Sanchez, M.-J. et al. An SCL 3’ enhancer targets developing endothelium together with embryonic and adult haematopoietic progenitors. Development 126, 3891–3904 (1999).

9. Sánchez, M.a.-J., Bockamp, E.-O., Miller, J., Gambardella, L. & Green, A.R. Selective rescue of early haematopoietic progenitors in Scl–/– mice by expressing Scl under the control of a stem cell enhancer. Development 128, 4815–4827 (2001).

10. Göttgens, B. et al. The scl +18/19 Stem Cell Enhancer Is Not Required for Hematopoiesis: Identification of a 5′ Bifunctional Hematopoietic-Endothelial Enhancer Bound by Fli-1 and Elf-1. Molecular and Cellular Biology 24, 1870–1883 (2004).

11. Liu, Y. et al. The genomic landscape of pediatric and young adult T-lineage acute lymphoblastic leukemia. Nature Genetics 49, 1211–1218 (2017).

12. Pölönen, P. et al. The genomic basis of childhood T-lineage acute lymphoblastic leukaemia. Nature 632, 1082–1091 (2024).

13. Begley, C.G. et al. Chromosomal translocation in a human leukemic stem-cell line disrupts the T-cell antigen receptor delta-chain diversity region and results in a previously unreported fusion transcript. Proceedings of the National Academy of Sciences 86, 2031–2035 (1989).

14. Brown, L. et al. Site-specific recombination of the tal-1 gene is a common occurrence in human T cell leukemia. Embo j 9, 3343–51 (1990).

15. Mansour, M.R. et al. An oncogenic super-enhancer formed through somatic mutation of a noncoding intergenic element. Science 346, 1373–1377 (2014).

16. Liu, Y. et al. Discovery of regulatory noncoding variants in individual cancer genomes by using cis-X. Nature Genetics 52, 811–818 (2020).

17. Smith, C. et al. TAL1 activation in T-cell acute lymphoblastic leukemia: a novel oncogenic 3’ neo-enhancer. Haematologica 108, 1259–1271 (2023).

18. Greig, K.T., Carotta, S. & Nutt, S.L. Critical roles for c-Myb in hematopoietic progenitor cells. Seminars in Immunology 20, 247–256 (2008).

19. Shao, X. et al. Transcriptional regulatory program controlled by MYB in T-cell acute lymphoblastic leukemia. Leukemia 38, 2573–2584 (2024).

20. Fuglerud, B.M., Ledsaak, M., Rogne, M., Eskeland, R. & Gabrielsen, O.S. The pioneer factor activity of c-Myb involves recruitment of p300 and induction of histone acetylation followed by acetylation-induced chromatin dissociation. Epigenetics & Chromatin 11, 35 (2018).

21. Lemma, R.B. et al. Chromatin occupancy and target genes of the haematopoietic master transcription factor MYB. Scientific Reports 11, 9008 (2021).

22. Sanda, T. et al. Core Transcriptional Regulatory Circuit Controlled by the TAL1 Complex in Human T Cell Acute Lymphoblastic Leukemia. Cancer Cell 22, 209–221 (2012).

23. Avsec, Ž., et al. Advancing regulatory variant eZect prediction with AlphaGenome. Nature 649, 1206–1218 (2026).

24. Sanda, T. & Leong, W.Z. TAL1 as a master oncogenic transcription factor in T-cell acute lymphoblastic leukemia. Experimental Hematology 53, 7–15 (2017).

25. Ong, J.Z.L., Tan, T.K., Wang, L., Tan, S.H. & Sanda, T. Regulatory mechanisms and context-dependent roles of TAL1 in T-cell acute lymphoblastic leukemia. Haematologica 109, 1359–1372 (2024).

26. Cordes, M. et al. Single-cell immune profiling reveals thymus-seeding populations, T cell commitment, and multilineage development in the human thymus. Science Immunology 7, eade0182 (2022).

27. Mansour, M.R. et al. The TAL1 complex targets the FBXW7 tumor suppressor by activating miR-223 in human T cell acute lymphoblastic leukemia. Journal of Experimental Medicine 210, 1545–1557 (2013).

28. Corces, M.R. et al. Lineage-specific and single-cell chromatin accessibility charts human hematopoiesis and leukemia evolution. Nature Genetics 48, 1193–1203 (2016).

29. Abraham, B.J. et al. Small genomic insertions form enhancers that misregulate oncogenes. Nature Communications 8, 14385 (2017).

30. Bernard, O. et al. A third tal-1 promoter is specifically used in human T cell leukemias. Journal of Experimental Medicine 176, 919–925 (1992).

31. Sharma, A. et al. Isoforms of the TAL1 transcription factor have diZerent roles in hematopoiesis and cell growth. PLOS Biology 21, e3002175 (2023).

32. Costa, J.R. et al. Transcription factor cooperativity at a GATA3 tandem DNA sequence determines oncogenic enhancer-mediated activation. Cell Reports 44, 115705 (2025).

33. Li, Y. et al. Alteration of CTCF-associated chromatin neighborhood inhibits TAL1-driven oncogenic transcription program and leukemogenesis. Nucleic Acids Research 48, 3119–3133 (2020).

34. Mifsud, B. et al. Mapping long-range promoter contacts in human cells with high-resolution capture Hi-C. Nature Genetics 47, 598–606 (2015).

35. Tan, T.K., Zhang, C. & Sanda, T. Oncogenic transcriptional program driven by TAL1 in T-cell acute lymphoblastic leukemia. International Journal of Hematology 109, 5–17 (2019).

36. Welcker, M. & Clurman, B.E. FBW7 ubiquitin ligase: a tumour suppressor at the crossroads of cell division, growth and diZerentiation. Nature Reviews Cancer 8, 83–93 (2008).

37. Bender, T.P., Kremer, C.S., Kraus, M., Buch, T. & Rajewsky, K. Critical functions for c-Myb at three checkpoints during thymocyte development. Nature Immunology 5, 721–729 (2004).

38. Stergiou, I.E. et al. EPH/Ephrin Signaling in Normal Hematopoiesis and Hematologic Malignancies: Deciphering Their Intricate Role and Unraveling Possible New Therapeutic Targets. Cancers 15, 3963 (2023).

39. Chen, J., Song, W. & Amato, K. Eph receptor tyrosine kinases in cancer stem cells. Cytokine & Growth Factor Reviews 26, 1–6 (2015).

40. Hoang, T., Lambert, J.A. & Martin, R. Chapter Six - SCL/TAL1 in Hematopoiesis and Cellular Reprogramming. in Current Topics in Developmental Biology, Vol. 118 (ed. Bresnick, E.H.) 163–204 (Academic Press, 2016).

41. Lausen, J. et al. Targets of the Tal1 Transcription Factor in Erythrocytes: E2 ubiquitin conjugase regulation by Tal1. Journal of Biological Chemistry 285, 5338–5346 (2010).

42. Murre, C. Helix-loop-helix proteins and lymphocyte development. Nature Immunology 6, 1079–1086 (2005).

43. Silberstein, L. et al. Transgenic Analysis of the Stem Cell Leukemia +19 Stem Cell Enhancer in Adult and Embryonic Hematopoietic and Endothelial Cells. Stem Cells 23, 1378–1388 (2005).

44. Huang, D. & Ovcharenko, I. The contribution of silencer variants to human diseases. Genome Biology 25, 184 (2024).

45. Dunning, A.M. et al. Breast cancer risk variants at 6q25 display diZerent phenotype associations and regulate ESR1, RMND1 and CCDC170. Nature Genetics 48, 374–386 (2016).

46. Huang, D. et al. Super-silencers are crucial for development and carcinogenesis in B cells. Nature Communications 16, 8395 (2025).

47. Mudge, Jonathan M. et al. GENCODE 2025: reference gene annotation for human and mouse. Nucleic Acids Research 53, D966–D975 (2024).

48. Flasch, D.A. et al. Somatic LINE-1 promoter acquisition drives oncogenic FOXR2 activation in pediatric brain tumor. Acta Neuropathologica 143, 605–607 (2022).

49. Zhang, J. et al. Deregulation of DUX4 and ERG in acute lymphoblastic leukemia. Nature Genetics 48, 1481–1489 (2016).

50. Buenrostro, J.D., Wu, B., Chang, H.Y. & Greenleaf, W.J. ATAC-seq: A Method for Assaying Chromatin Accessibility Genome-Wide. Current Protocols in Molecular Biology 109, 21.29.1–21.29.9 (2015).

51. Rao, Suhas S.P. et al. A 3D Map of the Human Genome at Kilobase Resolution Reveals Principles of Chromatin Looping. Cell 159, 1665–1680 (2014).

52. Kent, W.J., Zweig, A.S., Barber, G., Hinrichs, A.S. & Karolchik, D. BigWig and BigBed: enabling browsing of large distributed datasets. Bioinformatics 26, 2204–2207 (2010).

53. Li, H. & Durbin, R. Fast and accurate short read alignment with Burrows–Wheeler transform. Bioinformatics 25, 1754–1760 (2009).

54. Edmonson, M.N. et al. Bambino: a variant detector and alignment viewer for next-generation sequencing data in the SAM/BAM format. Bioinformatics 27, 865–866 (2011).

55. Chen, X. et al. CONSERTING: integrating copy-number analysis with structural-variation detection. Nature Methods 12, 527–530 (2015).

56. Hao, Y. et al. Dictionary learning for integrative, multimodal and scalable single-cell analysis. Nature Biotechnology 42, 293–304 (2024).

57. Rauluseviciute, I. et al. JASPAR 2024: 20th anniversary of the open-access database of transcription factor binding profiles. Nucleic Acids Research 52, D174–D182 (2023).

58. Kulakovskiy, I.V. et al. HOCOMOCO: towards a complete collection of transcription factor binding models for human and mouse via large-scale ChIP-Seq analysis. Nucleic Acids Research 46, D252–D259 (2017).

59. Bailey, T.L., Johnson, J., Grant, C.E. & Noble, W.S. The MEME Suite. Nucleic Acids Research 43, W39–W49 (2015).

60. Tanaka, H. et al. Interaction of the pioneer transcription factor GATA3 with nucleosomes. Nature Communications 11, 4136 (2020).

61. Cicirò, Y. & Sala, A. MYB oncoproteins: emerging players and potential therapeutic targets in human cancer. Oncogenesis 10, 19 (2021).

62. Bowers, S.R., Calero-Nieto, F.J., Valeaux, S., Fernandez-Fuentes, N. & Cockerill, P.N. Runx1 binds as a dimeric complex to overlapping Runx1 sites within a palindromic element in the human GM-CSF enhancer. Nucleic Acids Research 38, 6124–6134 (2010).

63. Gould, K.A. & Bresnick, E.H. Sequence determinants of DNA binding by the hematopoietic helix-loop-helix transcription factor TAL1: importance of sequences flanking the E-box core. Gene expression 7 **2**, 87–101 (1998).

64. Kharchenko, P.V., Tolstorukov, M.Y. & Park, P.J. Design and analysis of ChIP-seq experiments for DNA-binding proteins. Nature Biotechnology 26, 1351–1359 (2008).

65. Tischler, G. & Leonard, S. biobambam: tools for read pair collation based algorithms on BAM files. Source Code for Biology and Medicine 9, 13 (2014).

66. Li, H. et al. The Sequence Alignment/Map format and SAMtools. Bioinformatics 25, 2078–2079 (2009).

67. Zhang, Y. et al. Model-based Analysis of ChIP-Seq (MACS). Genome Biology 9, R137 (2008).

68. Ramírez, F. et al. deepTools2: a next generation web server for deep-sequencing data analysis. Nucleic Acids Research 44, W160–W165 (2016).

69. Hnisz, D. et al. Activation of proto-oncogenes by disruption of chromosome neighborhoods. Science 351, 1454–1458 (2016).

70. Langmead, B. & Salzberg, S.L. Fast gapped-read alignment with Bowtie 2. Nature Methods 9, 357–359 (2012).

71. Quinlan, A.R. & Hall, I.M. BEDTools: a flexible suite of utilities for comparing genomic features. Bioinformatics 26, 841–842 (2010).

72. Delaneau, O., Zagury, J.-F., Robinson, M.R., Marchini, J.L. & Dermitzakis, E.T. Accurate, scalable and integrative haplotype estimation. Nature Communications 10, 5436 (2019).

73. Hagiwara, K., Edmonson, M.N., Wheeler, D.A. & Zhang, J. indelPost: harmonizing ambiguities in simple and complex indel alignments. Bioinformatics 38, 549–551 (2021).

74. Durand, N.C. et al. Juicer Provides a One-Click System for Analyzing Loop-Resolution Hi-C Experiments. Cell Systems 3, 95–98 (2016).

75. Zhou, X. et al. Exploration of Coding and Non-coding Variants in Cancer Using GenomePaint. Cancer Cell 39, 83–95.e4 (2021).

76. Dobin, A. et al. STAR: ultrafast universal RNA-seq aligner. Bioinformatics 29, 15–21 (2012).

77. Anders, S., Pyl, P.T. & Huber, W. HTSeq—a Python framework to work with high-throughput sequencing data. Bioinformatics 31, 166–169 (2014).

78. Courtes, C. et al. Erythroid-specific Inhibition of the tal-1 Intragenic Promoter Is Due to Binding of a Repressor to a Novel Silencer. Journal of Biological Chemistry 275, 949–958 (2000).

